# A modular, immunopeptidogenomic (iPepGen) analysis pipeline for discovery, verification, and prioritization of cancer peptide neoantigen candidates

**DOI:** 10.1101/2025.04.07.647596

**Authors:** Subina Mehta, Reid Wagner, Katherine T. Do, James E. Johnson, Fengchao Yu, Tyler Jubenville, Kyle Richards, Suzanne Coleman, Flavia E. Popescu, Alexey I. Nesvizhskii, David A. Largaespada, Pratik D. Jagtap, Timothy J. Griffin

## Abstract

Characterizing tumor-specific neoantigen peptides, derived from genomic or transcriptomic aberrations and presented to the immune system, is critical for immuno-oncology studies. To this end, the modular iPepGen immunopeptidogenomics pipeline provides these functions: (1) Neoantigen prediction and protein database generation from genomic or transcriptomic sequencing data; (2) Peptide identification (3) Verification from immunopeptidomic mass spectral data; (4) Neoantigen classification and visualization; (5) Candidate prioritization for further study. Easy access via a publicly available, scalable cloud-based gateway coupled with online, interactive training materials streamlines the adoption by cancer researchers who require immunopeptidogenomic analysis tools but lack advanced computational expertise and resources.

## Background

Neoantigen peptides, derived from non-reference proteoforms uniquely expressed in tumor cells, are an increasingly important class of targets in cancer immunotherapy. These proteoforms may arise from a variety of genomic, transcriptomic, and post-translational mechanisms, including non-synonymous mutations, gene fusions [1], aberrant splicing, de novo transcription from non-coding regions [2], derepression of transposable elements, post-translational modification (PTM) [3], and translation of viral open reading frames in virus-associated cancers [4]. These alterations yield tumor-specific antigens absent from normal tissues and thus bypass central tolerance [5–8]. In addition to neoantigens, tumor-associated antigens (TAAs) and tumor-specific antigens (TSAs) derived from reference proteoforms whose expression is either enriched (TAAs) or only found in cancerous cells (TSAs) are of interest for immunotherapy development [9].

Once synthesized, aberrant proteoforms are degraded by cellular proteolytic machinery. In the cytosol, peptides of 8–11 amino acids are generated and presented by Major Histocompatibility complex (MHC) class I (called Human Leukocyte Antigen complex, HLA-A, -B, -C in humans) [10]. Meanwhile, longer peptides from extracellular proteins (10–25 amino acids) are presented via MHC class II (HLA-D family in humans) [11]. These complexes are displayed on the cell surface and recognized by T cells. It has been estimated that up to 10% of these peptides may elicit immunogenic responses when derived from non-reference sequences [12]. Despite this, tumors can escape immune surveillance via multiple mechanisms [13]. However, when MHC-bound neoantigen peptides are successfully identified, they can be leveraged for immunotherapy, including adoptive T cell transfer, immune checkpoint blockade, and prophylactic vaccination [14–18].

### Identifying and predicting neoantigen candidates from genomic and transcriptomic data

The ability to identify neoantigen candidates from genomic and transcriptomic sequences has advanced rapidly due to improvements in next-generation sequencing (NGS) and customized computational pipelines [8,19]. Numerous tools have been developed for identifying potential neoantigens encoded by tumor-specific alterations [20–24]. These tools typically utilize RNA-Seq data or whole-exome sequencing to genotype alleles coding for MHC protein subunits and predict binding affinities of potential neoantigen peptides to the MHC based on biochemical and structural modeling [25–28]. Higher predicted binding affinity to specific MHC (or HLA) alleles is an indicator of immune recognition and the possibility of a peptide to elicit an immunogenic response. Deep learning tools have also emerged that seek to predict potential immunogenicity for neoantigen sequences identified by genome or transcriptome sequencing [29–31].

### Beyond prediction: confirming MHC-bound neoantigens via mass spectrometry (MS)-based immunopeptidomics

Although important, NGS-based approaches ultimately provide only a predictive view of antigen presentation, which presents limitations [8,32]. These include the inability to confirm proteoform translation, peptide processing, or MHC presentation. Predictions also fail to capture PTMs, which are known to modulate immunogenicity [33], are prone to prediction bias [34,35], and are often missed as antigenic peptides [36]. Empirical follow-up studies have shown that only a small fraction of predicted neoantigens from NGS data are processed, bound to the MHC, and truly immunogenic [17]. These constraints have motivated the development of mass spectrometry (MS)-based immunopeptidomics approaches that allow direct identification of peptides presented by MHC complexes [37–39].

In immunopeptidomics, the intact MHC and bound peptides are affinity-enriched and analyzed by liquid chromatography (LC) tandem mass spectrometry (MS/MS). Tandem mass spectra are then matched to custom protein databases that include both canonical reference sequences and tumor-specific neoantigen candidates inferred from NGS data. This strategy allows the detection of neoantigens and TAAs/TSAs in an unbiased, high-throughput manner, including peptides with PTMs that might influence immunogenicity [33]. Originally developed by Hunt and colleagues [40], MS-based immunopeptidomics has matured and now enables the routine detection of thousands of MHC-bound peptides from cellular and tissue sources [39,41], significantly expanding the scope of immuno-oncology research [8,38].

### Informatics requirements for neoantigen prediction and confirmation via immunopeptidogenomics

The integration of NGS and MS data for comprehensive neoantigen discovery, appropriately called “immunopeptidogenomics” by Purcell and colleagues [42], poses unique informatics requirements and challenges. These include managing the heterogeneity of NGS alterations for predicting neoantigens and constructing sample-specific databases; efficient and sensitive peptide spectrum match (PSM) generation from MS/MS searched against customized protein sequence databases using non-specific enzyme constraints [33,43,44], and, if desired, analyzing PTMs; quantifying identified peptides via label-free methods [45,46].

Some pipelines have been described that focus on the analysis of MS-based immunopeptidomics data [47–49]. To varying degrees, these bioinformatic platforms offer the ability to identify potential genomic/transcriptomic aberrations from NGS data and utilize tools for predicting MHC binding to help in prioritization. Some of the platforms [47,48] use publicly available, established software such as MaxQuant [50] or commercial algorithms [42] for generating PSMs from MS/MS data; however, none of these algorithms are optimized for analysis of MHC-bound peptides using “no enzyme” settings, rather than peptides generated by trypsin digestion, as is common in proteomics applications. Recently, a sophisticated multi-omics platform called NeoDisc has been described that is openly available for non-commercial use [51]. NeoDisc offers a complete pipeline with modules for neoantigen prediction, peptide identification from immunopeptidomic MS/MS data, and advanced methods, including a deep learning algorithm for prioritizing candidates for further study. However, this platform requires substantial technical expertise for installation, as well as local computational infrastructure to support its complex set of tools and their computing requirements. It also lacks a GUI, limiting its accessibility and usage by non-expert users who may be new to immunopeptidogenomics. Moreover, the addition of new or experimental tools into these previously described pipelines requires significant software developer expertise, beyond the reach of most non-expert users.

### The iPepGen pipeline and the Galaxy ecosystem as a unique solution for immunopeptidogenomic informatics

To address current limitations, we have developed the modular immunopeptidogenomic (iPepGen) pipeline for the discovery and characterization of neoantigens. Deployment in the Galaxy ecosystem [52,53] provides a web-based environment and user interface, immediately accessible via cloud-hosted servers [54], that supports reproducible bioinformatics analyses. Tools are openly available via the Galaxy Tool Shed and can be deployed on public cloud servers or institutional high-performance computing (HPC) resources. Galaxy supports comprehensive NGS and MS-based proteomics workflows [55], with built-in provenance tracking, visualizations, and shareable software workflows and analysis histories. The powerful Galaxy Training Network (GTN) [56,57] further promotes community adoption by offering interactive training resources for new users. Leveraging these strengths, iPepGen integrates validated tools for neoantigen prediction, peptide identification and verification, genomic mapping, and immunogenicity assessment within a unified and accessible framework complete with training resources. Taken together, these features lower the entry bar for adoption by new users, especially those lacking advanced computational expertise and/or access to HPC resources. Overall, the iPepGen pipeline offers a unique informatics resource built to advance studies by a growing community of immuno-oncology researchers using immunopeptidogenomics for neoantigen discovery and characterization.

## Results

We implemented the iPepGen pipeline within the Galaxy ecosystem to support end-to-end neoantigen discovery and prioritization by integrative analysis of NGS and LC-MS/MS immunopeptidomics data, along with other specialized tools. To demonstrate the effectiveness and reproducibility of the modular pipeline, we analyzed paired RNA-Seq and HLA class I immunopeptidomics data from a human malignant peripheral nerve sheath tumor (MPNST) cell line, STS-26T. Moreover, to highlight the flexibility and effectiveness of iPepGen in addressing the challenges of immunopeptidogenomics analysis, we show results from analysis of additional multi-omics data available from previously published studies.

### A modular, cloud-based pipeline is easily accessed and used via a web-based interface

The iPepGen pipeline is composed of five modular workflows (**Figure 1**), deployed in the Galaxy bioinformatics environment. These workflows integrate software tools selected and optimized to accomplish specific data analysis tasks and generate results compatible with subsequent analysis by downstream modules in the pipeline. The workflows composing each module are freely available on the European Galaxy server, through the links shown below in **Table 1**. A Galaxy workflow contains all software tools for operation on appropriate input data, along with preset, optimized settings. We have made example input files available for each workflow via Galaxy links **(Table 1)**, which can be imported and recognized as appropriate input formats by the corresponding workflow. Once a workflow is run, Galaxy saves the analysis as a history, which contains all input and result files generated by the workflow. The history can be shared, re-run using adjusted settings, and/or archived. Links to completed histories for each iPepGen module are also shown in **Table 1**, using the example MPNST data. Workflows and histories can be imported into any Galaxy account once a user has registered on the server.

**Figure 1.**
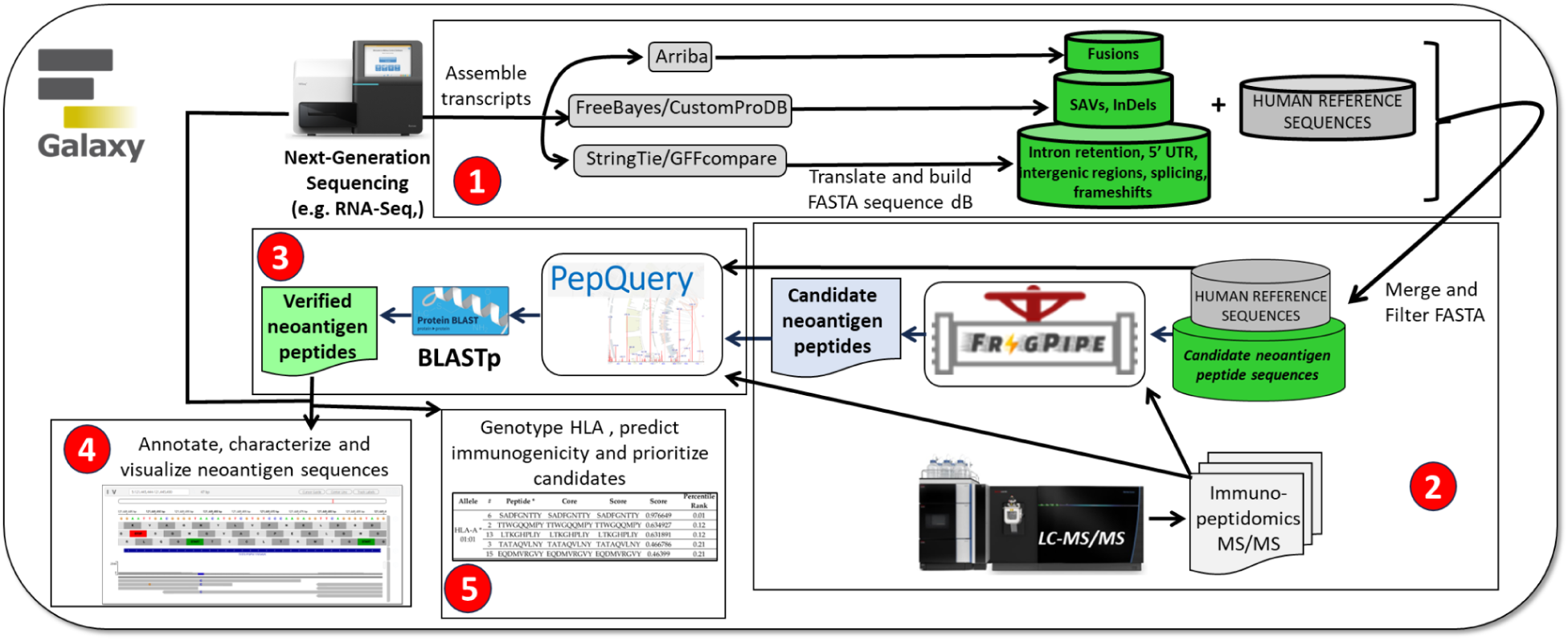
Five modules comprising the Galaxy-based iPepGen pipeline. The modules carry out the following functions: 1) Predict neoantigen candidates from NGS data and generate customized protein sequence databases, including reference and non-reference neoantigen candidate sequences; 2) Discover neoantigen peptide candidates by sequence database searching of tandem mass spectrometry (MS/MS) immunopeptidomics data; 3) Verify discovered peptide candidates through a secondary peptide-centric evaluation against the MS/MS dataset; 4) Visualize and classify the nature of verified neoantigen peptides encoded by the genome and/or transcriptome; 5) Prioritize neoantigens for further exploration and empirical validation.

**Table 1.**
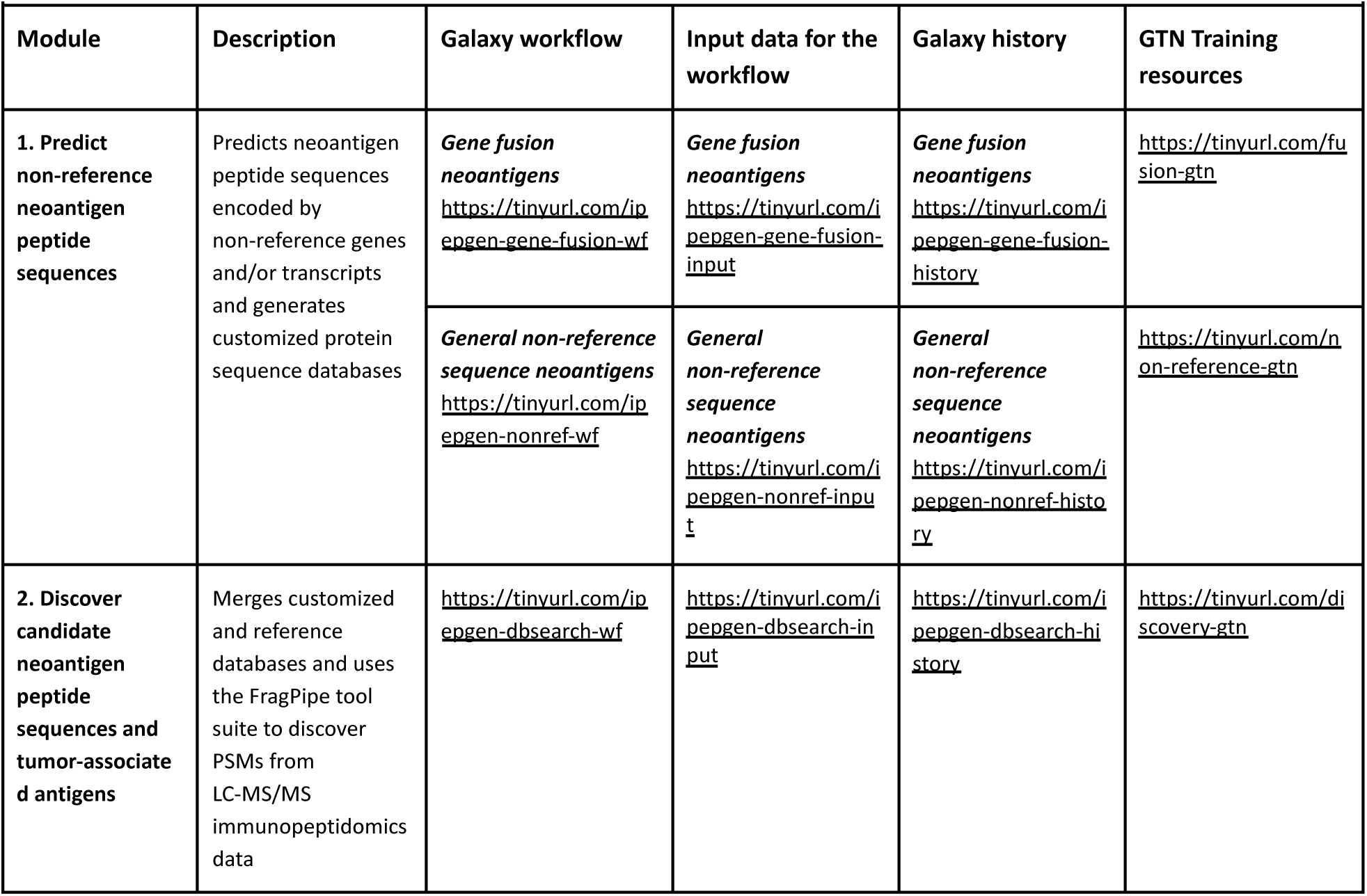
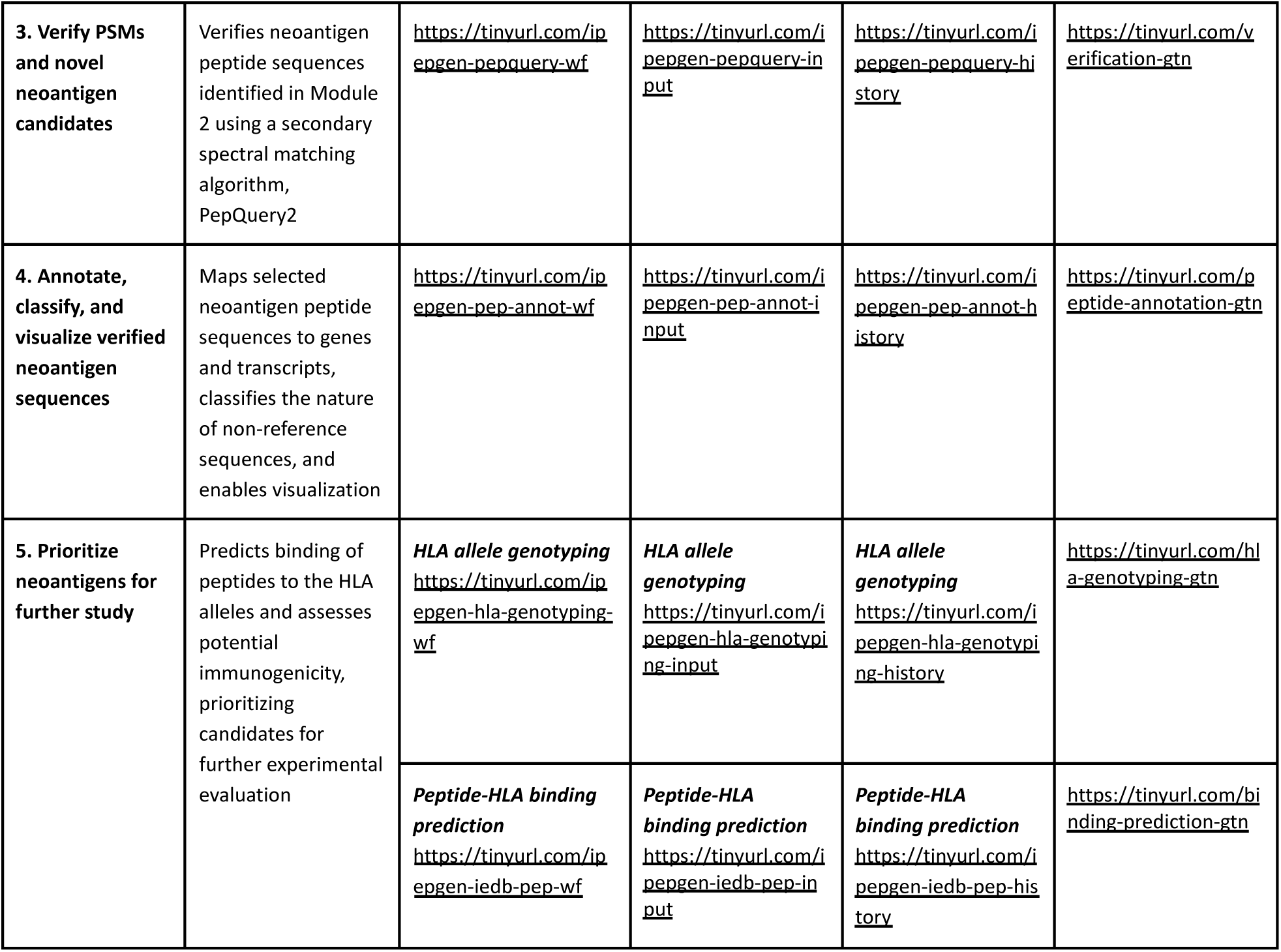
Links for workflows, input data, analysis histories, and training resources for each module of the iPepGen analysis pipeline.

### Online, on-demand, and interactive training resources lower barriers for new users

The iPepGen pipeline takes advantage of the powerful GTN [56], which offers online, on-demand, and interactive training resources for new users. The GTN is seamlessly connected with workflows and software available on public Galaxy servers, thus providing a means for new users to adopt sophisticated analysis pipelines for use in their studies. We developed an accessible “Learning Pathway” [58] within the GTN, which summarizes training resources for each module in the iPepGen pipeline, and leads users to the online documentation. The last column of **Table 1** shows the links to training materials that guide new users step-by-step through each workflow, complete with links to demonstration data and software. Once a user has completed these training workflows, they are equipped with the necessary knowledge to upload their data to their Galaxy account and begin their own meaningful immunopeptidogenomic analysis.

### Description of analysis modules and representative results

We describe the main functionalities and operation of each module in the iPepGen pipeline, along with results from MPNST data that demonstrate its effectiveness for immunopeptidogenomic analysis. The Methods section provides a more detailed description of software tools and key settings used in each module.

### Module 1: Prediction of non-reference neoantigen peptide sequences

To demonstrate the functionality and output of the iPepGen pipeline, we analyzed RNA-Seq and immunopeptidomics data from an MPNST cell line. These demonstration analyses used the workflows and input data shown in **Table 1** and produced the analysis histories shown in this table as well. For Module 1, we utilize several complementary workflows seeking to cast a wide net for possible genomic or transcriptomic sequences that may code for novel proteoforms leading to neoantigen production and may be identified via matching to immunopeptidomic MS/MS data in downstream modules.

#### Gene Fusion Variant Detection

The first workflow targets gene fusions, a potential source of neoantigen peptides in many cancers [1], using the Arriba algorithm [59], integrated with the RNA STAR aligner [60] for spliced alignment and chimeric read detection. Outputs include a tabulated list of high-confidence fusions with supporting read metrics, visual summaries of each event in PDF format, and translated fusion junction sequences in FASTA format for downstream use (**Table 1**, Module 1, Gene Fusion neoantigens Item 21 in the history). Post-processing steps are applied to remove potential false positives based on read depth, mapping quality, and known benign fusion sequences. For the MPNST cell line RNA-Seq data used for demonstration, we identified 86 novel fusion sequences that were translated into potential neoantigen peptide sequences and outputted in the annotated FASTA-formatted table (**Figure 2**).

**Figure 2.**
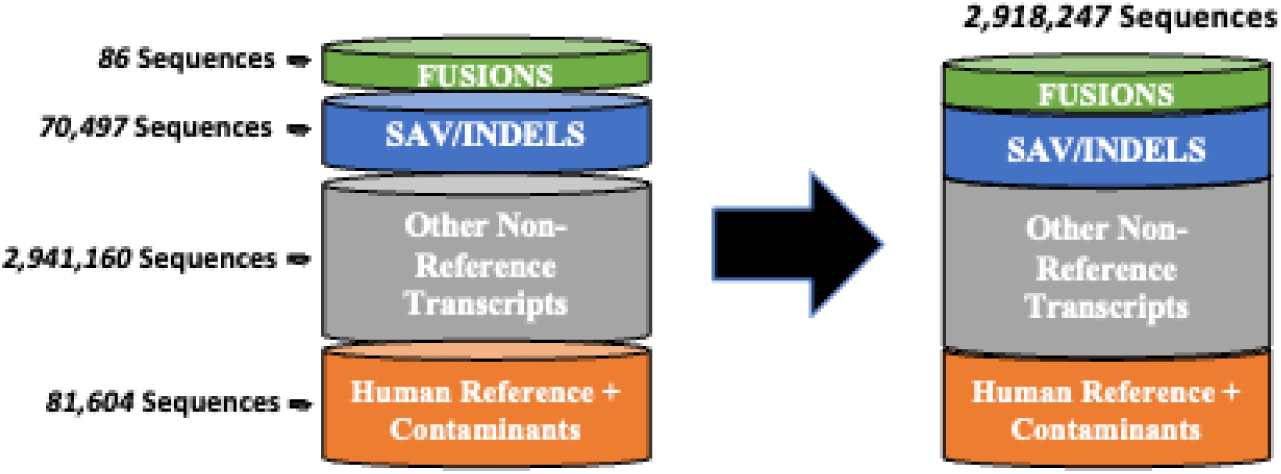
Relative size of databases from MPNST cell line RNA-Seq data generated by Arriba (fusions), CustomProDB (SAV/Indels) and StringTie (other non-reference sequences) workflows. The graphic illustrates the cumulative size of each non-reference database component and the final customized database after merging with the reference and common contaminant (cRAP) protein sequences and after removing redundant sequence entries.

#### Comprehensive Prediction of Non-Reference Protein Sequences

Complementing the targeted prediction of gene fusion-derived neoantigens, we employed additional workflows that broadly identify non-reference transcript sequences from RNA-Seq data. In one of these workflows, paired-end reads are aligned to the human reference genome using HISAT2 [61], followed by variant calling with FreeBayes. CustomProDB [62] then annotates predicted coding DNA sequence (CDS) variants, generating a set of non-reference protein sequences containing single amino acid variants (SAVs) and short insertion-deletions (InDels).

In the second workflow, transcript assembly is performed using StringTie [63], with transcript classification via GffCompare [64] to distinguish novel sequences compared to the reference genome. Novel, non-reference sequences, including frameshifts, novel splice events, and/or sequences from non-coding genomic regions, are translated in all three reading frames using Galaxy tools “Translate BED Transcripts” and “bed to protein map.”

The peptide sequences predicted from fusions, SAVs/InDels, and other non-reference transcripts are all merged with UniProt canonical human protein sequences and common contaminant sequences (called the common Repository of Adventitious Proteins, cRAP) using “FASTA Merge Files and Filter Unique Sequences.” Non-reference entries are annotated with a “Generic” header, distinguishing them from UniProt canonical sequences, to support downstream analysis in Module 2. For the RNA-Seq from the MPNST STS-26T cell line, these combined workflows generated a protein database containing approximately 2.9 million unique protein sequences (**Table 1**, Module 1, Item 41 in history). **Figure 2** shows the FASTA-formatted databases generated by each of the three described workflows used for the prediction of neoantigen peptides in this module using this MPNST demonstration data. Notably, the cumulative number of entries across all individual databases exceeds that of the comprehensive database, as redundant or overlapping peptide sequences present in multiple workflow-specific outputs are collapsed into unique entries during the database merging process, thereby reducing redundancy in the final merged FASTA database.

### Module 2: Discovery of candidate neoantigen peptide sequences and tumor-associated antigens

To identify candidate neoantigens presented by HLA class I molecules, we analyzed MS/MS immunopeptidomics data from the same MPNST cell line used in Module 1. This module utilizes the FragPipe software suite [65], which is deployed as a tool in Galaxy. We matched MS/MS spectra against the customized FASTA database of ∼2.9 million sequences generated previously (**Figure 2**). The workflow integrates MSFragger for PSM generation, with MSBooster rescoring [66], Percolator [67] for post-search validation, and Philosopher [68] for downstream filtering and false discovery rate (FDR) control at 1% for both PSM and protein levels. FragPipe also offers the IonQuant tool [46] for label-free quantification (LFQ), quantifying identified peptide abundance based on their normalized LC-MS intensities. The “Nonspecific-HLA” search mode in FragPipe is used to identify HLA I-bound peptides ranging from 8 to 20 amino acids in length. To accommodate the large search space when using no enzyme specificity, the workflow automatically partitions the protein database into 200 random sections for efficient parallel processing. For more efficient use of FragPipe in iPepGen, we developed a Galaxy tool called “Manifest Generator” for generating the tabular manifest experimental design file, which annotates input raw MS/MS scan files being analyzed into experimental groups defined by the user (e.g., control, treated). This file is required by FragPipe for operation and provides a comparison of LFQ values of identified peptides across sample groups, presented in the tabular output of all identified peptides from the analysis.

After the FragPipe analysis is complete, the Query Tabular Galaxy tool [69] extracts peptide sequences from the tabular outputs of FragPipe that did not match any canonical reference proteins, generating a new tabular file of candidate neoantigen peptides for further analysis. Users may further stratify these peptides by variant type (e.g., SAVs, InDels, gene fusions, etc.) by filtering on tag-based sequence annotations embedded during FASTA construction in Module 1. Using a single MS/MS file from the MPNST cell line STS-26T, and the comprehensive sequence database generated in Module 1, we identified 123 unique peptide sequences that mapped exclusively to non-reference proteins (**Table 1**, Module 2, Item 21 in history).

One key aspect of immunopeptidomics is the requirement to identify peptides using non-specific enzyme settings, as opposed to traditional bottom-up proteomics, which is based on analysis of peptides generated using proteolysis with trypsin. With non-specific enzyme settings, the possible sequence search space increases significantly, especially when using large databases including non-reference protein sequences predicted from NGS data. This challenges sequence database search algorithms to sensitively and efficiently generate PSMs for neoantigen discovery. FragPipe offers an ultrafast algorithm, which, when coupled with deployment in Galaxy on scalable compute resources, overcomes these limitations. As a demonstration, we compared FragPipe with the popular MaxQuant software, used in some available immunopeptidomics pipelines [50], evaluating their ability to identify HLA 1-bound peptides using non-specific enzyme settings (**Figure 3A**). Both tools were run on equivalent compute environments (16 cores, 64 GB memory). We limited the size of the customized protein sequence database to 250,000 entries because MaxQuant would not reliably run with larger databases, further demonstrating its limitations for neoantigen discovery. FragPipe identified a greater number of peptides at a global FDR of 1% while requiring significantly less runtime. This performance advantage can be attributed to MSFragger’s ultrafast search capabilities, including database splitting, and enhanced scoring strategies and post-processing methods [70], making it particularly well-suited for high-throughput neoantigen discovery. We also benchmarked the performance of FragPipe based on the size of the input sequence database. Unsurprisingly, the analysis time and memory required scale with the size of the database, with FragPipe generating results on even the very large 2.9M sequence database in under two hours on the publicly available European Galaxy server (see **Supplemental Table 1**).

**Figure 3.**
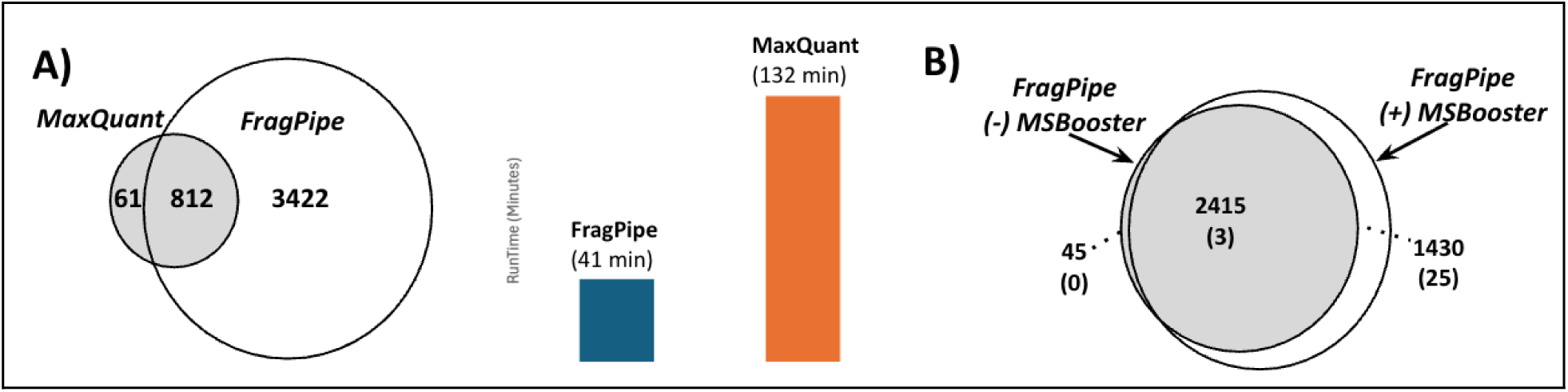
**A.** Comparison of FragPipe and MaxQuant performance for a “no enzyme” database search of a single peptidomics MS/MS file against an immunopeptidomics proteome database of 250K sequences from MPNST data in Module 1; both software were run using their stand-alone GUI using 16 cores and 64GB memory. **B.** Comparison of FragPipe run without and with MSBooster functionality on a single immunopeptidomics MS/MS datafile from MPNST cells and searched against a sequence database of 2.9M+ sequences, including the reference and annotated non-reference candidate neoantigen sequences; number of unique peptides identified are shown, with numbers in parentheses () indicating the number of candidate neoantigen peptides identified with sequences that do not map to the reference proteome.

In addition to MaxQuant, we also compared FragPipe’s performance for no enzyme PSM generation from large sequence databases to the commercial software PEAKS, an efficient algorithm for PSM generation that has also been used in previously described immunopeptidomics pipelines [42]. FragPipe identified 30% more peptides than PEAKS for database searching against our large demonstration database containing over 2.9M sequences (See **Supplemental Figure 1**).

We also evaluated the effects of using the MSBooster algorithm [66] in FragPipe, which utilizes machine-learning-based prediction of fragment ion intensities and retention times, followed by a re-scoring of potential PSMs to increase confident peptide identifications. **Figure 3B** shows the use of MSBooster on our example immunopeptidomics dataset from MPNST cells, generating PSMs against the large protein sequence database. MSBooster significantly increases the number of confident PSMs generated, including identification of candidate neoantigen sequences. We have also tested the use of MSBooster on other datasets, where it showed an increase in total peptides identified of about 35% on average, and an increase of about 50% in identified non-reference sequences (**Supplemental Figure 2**).

### Module 3: Verification of neoantigen peptide-spectrum matches

To further evaluate candidate neoantigens identified in Module 2, we implemented a secondary peptide-centric verification step using PepQuery2 [71]. This approach addresses the potential for false-positive PSMs when searching large, non-reference databases, as previously noted in immunopeptidomics studies [72]. PepQuery2 allows for independent evaluation of individual PSMs by scoring candidate neoantigen sequences against MS/MS spectra in the presence of competing reference peptides, including common PTMs. This module also uses the BLAST-P tool [73] for verifying that neoantigen candidates are truly novel and do not match any known reference sequences. As such, the module provides a secondary verification of candidate neoantigens identified using FragPipe against the large sequence database to ensure users are provided the most confident and high-value peptides to consider for further study.

For the demonstration MPNST immunopeptidomics dataset, we evaluated the 123 candidate neoantigen peptides from Module 2 using PepQuery2. Nineteen neoantigen peptide sequences passed PepQuery2 hyperscore validation and BLAST-P filtering (**Table 1**, Module 3, Item 12 in history). BLAST-P alignment was performed at less than 100% sequence identity to canonical proteins to capture sequence variants and polymorphisms. These peptides represent the most confident neoantigen candidates ready for further characterization and prioritization for experimental evaluation. We include **Supplemental Figure 3,** which shows the multiple steps our pipeline utilizes in order to discover neoantigen candidates and verify those of highest confidence, generating a stringently filtered set of peptides for further assessment.

### Module 4: Visualization and classification of verified neoantigen sequences

This module facilitates the classification and genomic annotation of verified neoantigen peptides. The PepPointer tool [74] is used to map neoantigen peptides to their genomic coordinates using BED and GTF annotations and classify their genomic origin as coding (CDS), untranslated region (UTR), intron retaining, or intergenic sequences. PepPointer outputs a tabular file with a description of their likely genomic origin, as well as genomic coordinates to use with the Integrated Genome Viewer (IGV) [75] and a URL for direct input into the UCSC Genome Browser [76], enabling researchers to inspect peptide-transcript-genome alignments visually. **Figure 4** shows an example output of a neoantigen candidate peptide from MPNST, visualized in IGV and mapped to the genome along with supporting assembled transcript sequences, generated in Module 1, that led to the predicted sequence that was verified in Modules 2 and 3.

**Figure 4.**
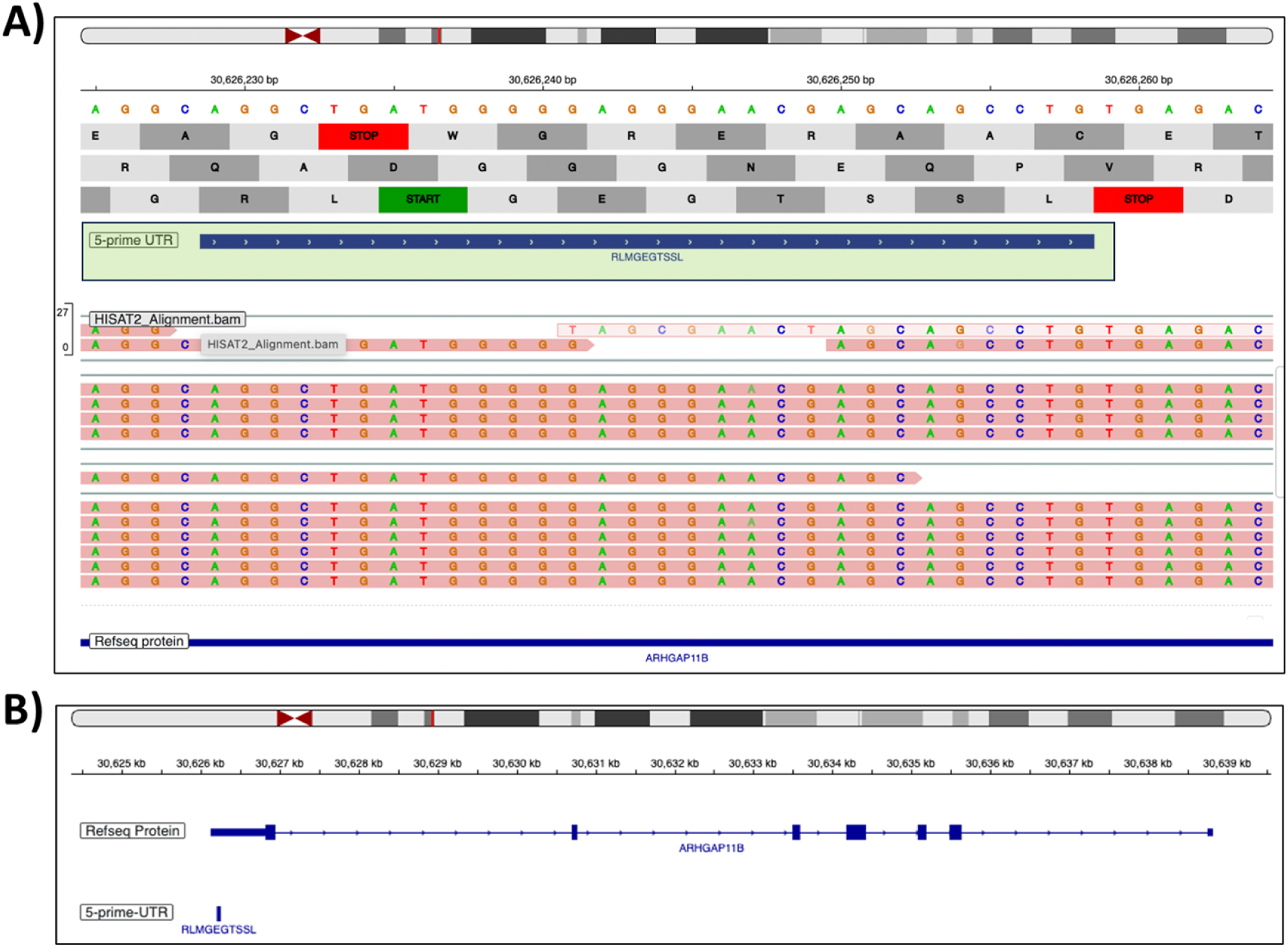
Example of a non-reference neoantigen peptide sequence (RLMGEGTSSL, shown in green shaded box) coded by a transcript sequence mapped to a portion of the 5’ UTR of an annotated gene structure and visualized in the Integrated Genome Viewer (IGV). Assembled RNA-Seq reads coding this peptide are in pink; B) A larger view of the annotated protein coded by this genomic location (exon coding sequences shown in blocked blue) with the location of the novel neoantigen peptide mapped to the 5’ UTR also shown as a reference.

Visualization in the UCSC browser using the URL generated by PepPointer offers a variety of tools for assessing the nature of neoantigen candidates and their potential for immunotherapy development. Visualization tracks can be easily added to assess gene expression in normal and cancerous tissues from the Genotype-Tissue Expression (GTEx) database [77], as well as gene expression profiles in cancerous tissues through resources such as Clinical Interpretation of Variants in Cancer (CIViC) [78] and the Cancer Genome Atlas (TCGA) cancer expression database [79]. **Supplemental Figure 4** shows an example of the same verified neoantigen peptide candidate shown in **Figure 4**, but visualized in the UCSC browser using some of these functionalities. Moreover, the tabular output provides users an easy means to input verified neoantigen peptide sequences into other resources, such as the PeptideAtlas resource [80], to determine if a peptide has been identified in other published MS-based peptidomics studies, or other repositories of neoantigens identified in prior, published studies [81,82].

In some cases, candidate neoantigen peptides arise from transcript coding sequences derived from complex processing of DNA or RNA sequences (e.g., unique combinations of intron and exon sequences), which are not easily annotated via PepPointer and may require manual exploration by the user for their advanced characterization. **Supplemental Figure 5** outlines one such example found in the nineteen neoantigen candidates outputted in Module 4 for the MPNST demonstration data (peptide HSEVQTLKY). Although this mapped to an intronic region, it required more manual investigation. **Table 2** summarizes the panel of nineteen verified neoantigen peptides from the demonstration data. These peptides were supported by high FragPipe PSM score probabilities (≥0.95) and deemed confident by PepQuery2 verification with highly significant p-values (≤1.0E-03). Notably, peptides in **Table 2** were derived from both canonical (CDS, UTR) and non-canonical genomic coding regions (introns, intergenic), underscoring the potential of iPepGen to uncover promising neoantigen peptides from diverse genomic and/or transcriptomic aberrations.

**Table 2.**
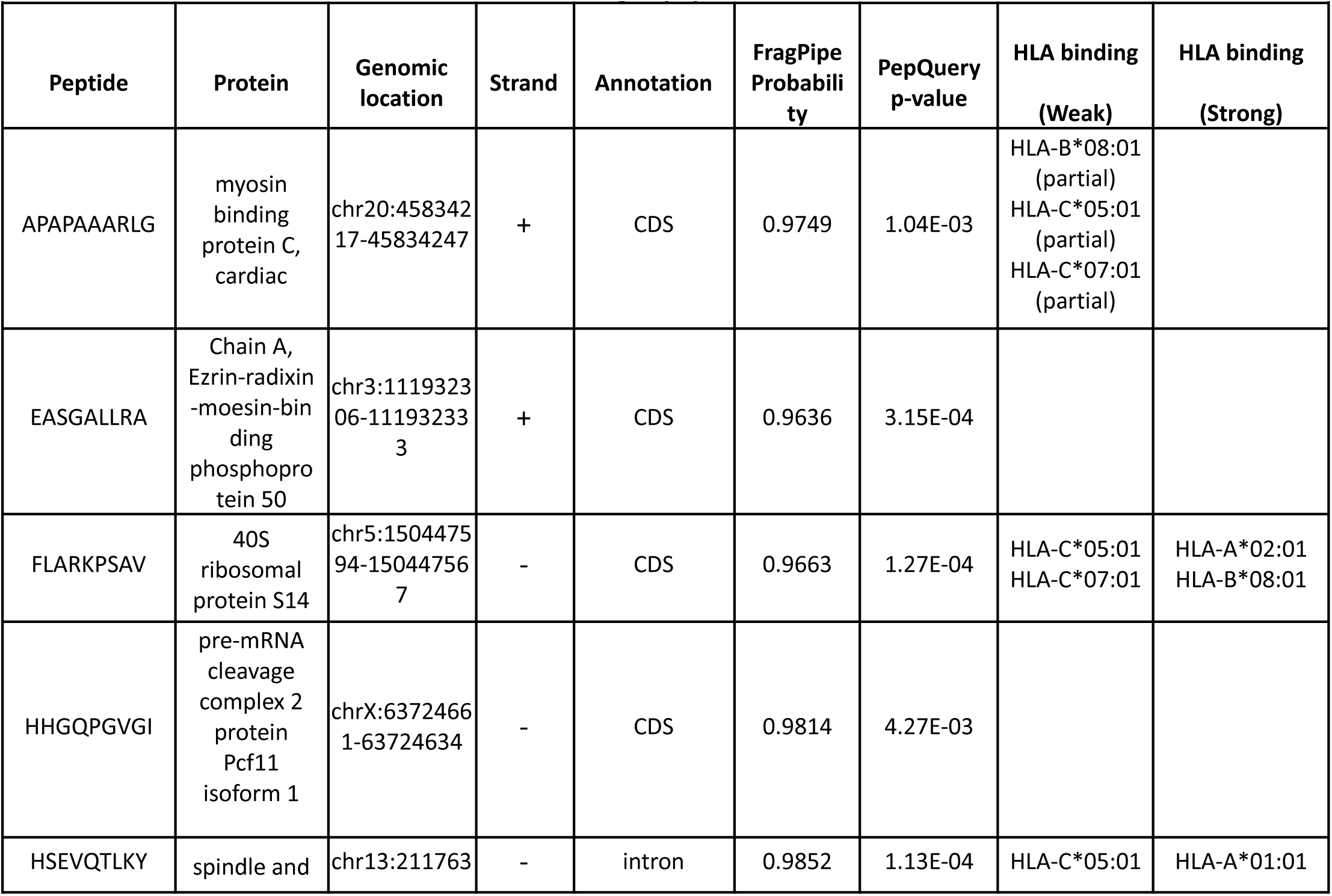
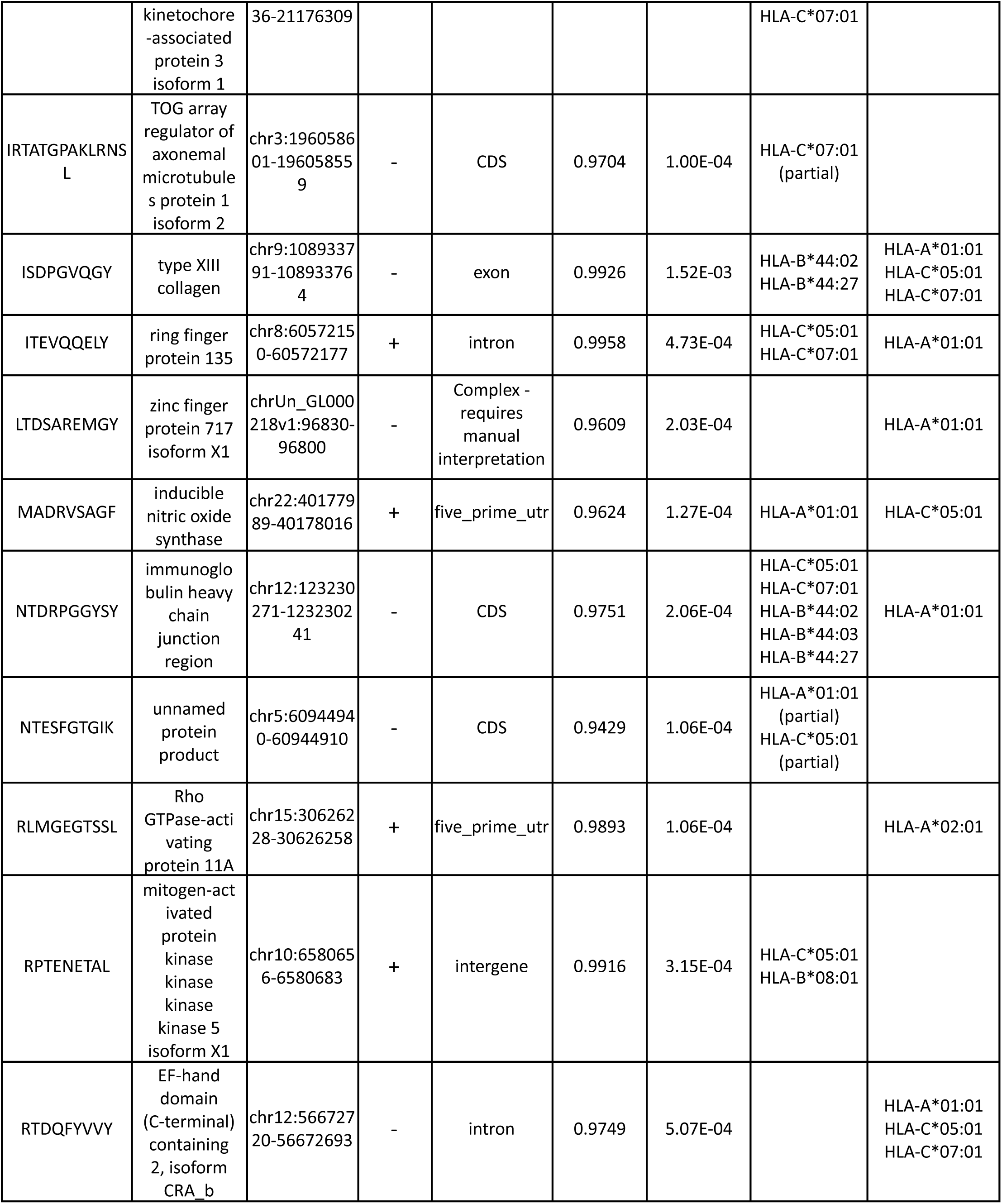
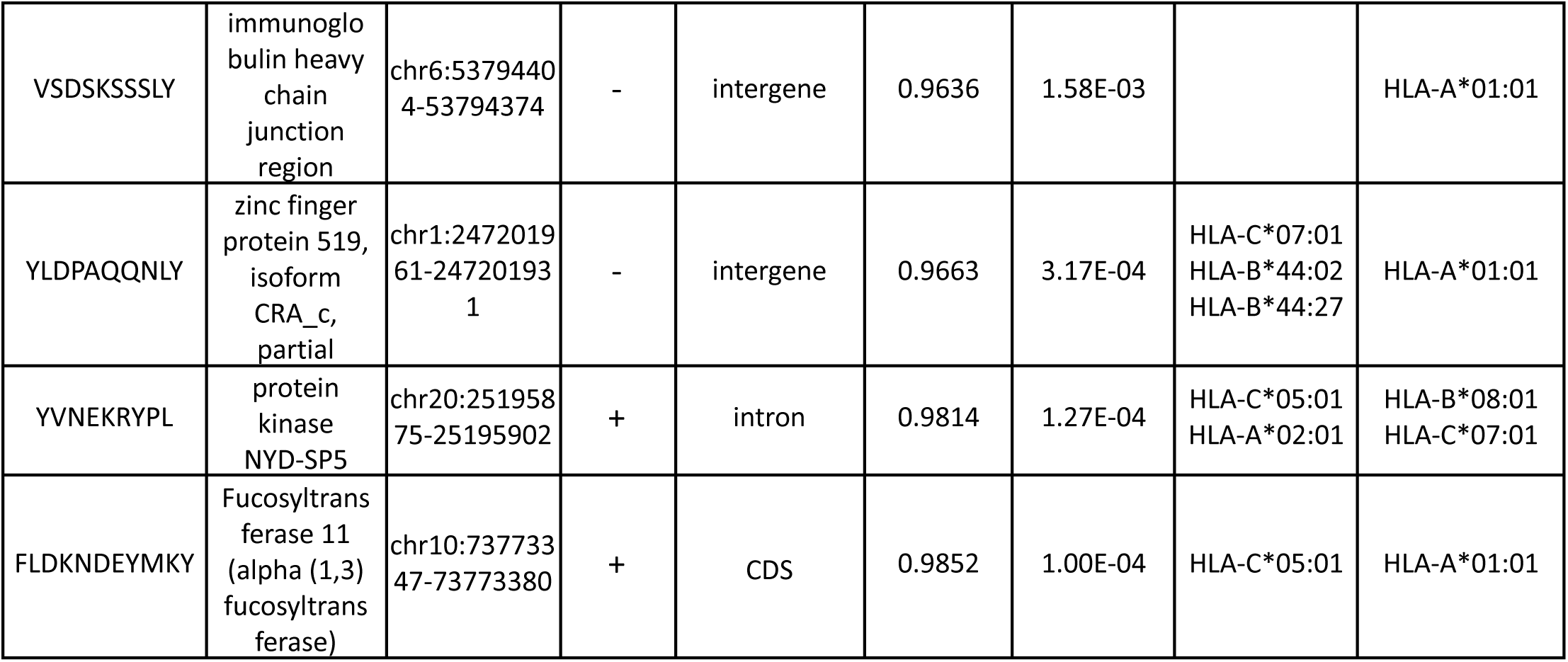
Verified and classified neoantigen peptides from MPNST demonstration data.

### Module 5: Prioritization of neoantigens for further study

With verified neoantigen peptides in hand from the modules above, iPepGen offers a module to assist users in prioritizing those sequences that may be of highest interest for further evaluation in the lab. This prioritization is guided using software for predicting binding and potential recognition as immunogenic epitopes that may be of highest value for immunotherapies. Two workflows make up this module to enable this prioritization.

#### HLA Genotyping

As a first step to predicting peptide binding to the MHC (or HLA for human samples), allele genotyping is performed on RNA-seq data. Paired-end reads in FASTQ format are analyzed using two complementary tools: OptiType [83], optimized for class I alleles, and Seq2HLA [84], which also covers class II loci, if desired. Outputs from the two tools are merged, and redundant entries are removed using Galaxy text manipulation tools. In the case of MPNST RNA-Seq data used for demonstration purposes, the final genotype set included eight HLA class I alleles (**Table 1**, Module 5, Item 13 in first history), which served as input for further neoantigen peptide-binding affinity prediction.

#### Peptide–HLA Binding Prediction

The genotyping results are used as one input to the IEDB MHC Binding Prediction tool [85], using the NetMHCpan_EL algorithm, trained on empirical immunopeptidomics data. The neoantigen peptides from Module 3 are used as the other input. If the input peptides are 9 or more amino acids long, the software generates overlapping shorter sequences of at least 8 amino acids and evaluates each for HLA binding from human samples, thereby expanding the list of potential peptide binders beyond the original input set. Peptide sequences are assigned a percentile rank for their binding to each HLA allele and classified as strong (percentile rank ≤ 0.5%) or weak binders (0.5 – 2%). Binding results for the neoantigen candidates identified from the MPNST dataset are shown in **Table 2**. Thirteen peptides were predicted as strong binders (**Table 1**, Module 5, Item 12 in the second history), and ten were classified as weak binders (**Table 1**, Module 5, Item 11 in the second history) to specific HLA alleles. The specific combination of predicted binding affinity to specific HLA alleles can be used as an indicator of immune recognition [86] and provides a means to prioritize peptides for further experimental evaluation.

#### Enhanced prioritization and immunogenicity prediction

The outputted HLA allele genotypes and verified neoantigen candidate peptide sequences from iPepGen serve as input to the pVACbind software suite, which we deployed as a Galaxy tool. pVACbind is part of the personalized Variant Antigens in Cancer tool suite (pVACtools) [24]. The pVACbind software bundles numerous complementary tools for predicting binding of input peptide sequences to HLA protein complexes (both class I and II) coded by specific allele genotypes, as well as two deep learning algorithms, DeepImmuno [29] and BigMHC [30], for predicting immunogenicity of HLA alleles and bound peptides. The Galaxy pVACbind tool outputs a tabular file with aggregated results across all the algorithms selected by the user, as well as a filtered output showing only those peptide and allele combination scores indicating high potential for immunogenicity. The pVACbind software is well documented [87] and provides users with ample information about the bundled tools and how to interpret results for assisting in prioritizing peptides for further experimental testing as immunotherapy agents. We have documented the use of pVACbind in Galaxy in the GTN, and also in **Supplemental Figure 6 (a, b, c)**.

### Extended applications of the modular iPepGen pipeline

#### An end-to-end “One-Click” iPepGen workflow

We have intentionally used a modular design in developing the iPepGen pipeline, as described above in detail. This design provides the best means to develop module-specific online training material accessible through the GTN, aimed at training new users seeking to adopt these tools and apply them to their own data. However, for more advanced users who have trained through the GTN, a module-by-module analysis may be cumbersome, especially for studies generating multiple MS-based immunopeptidomics datasets requiring reproducible analysis [88]. To this end, we developed a comprehensive workflow integrating all the modules of iPepGen. In this embodiment, the user simply uploads their starting raw data (e.g., paired RNA-Seq, LC-MS/MS scan files) along with the required reference proteome sequence database, genome sequence annotation files, and manifest experimental design file. The entire pipeline is instantiated with a single click of the “Run” button for the workflow. Although all intermediate files are recorded and saved in the resulting Galaxy analysis history, we have simplified the outputs of this integrated workflow so only those of the highest value for interpreting results are exposed to the user in the generated history.

We developed the one-click workflow using our single MPNST dataset. To further demonstrate its effectiveness, we utilized publicly available data from a recent immunopeptidogenomic study by the Purcell group [89]. RNA-Seq data from an acute myeloid leukemia cell line were selected along with three raw MS/MS files of technical replicates from an experiment testing an HLA class I immunopeptidomic enrichment method. Links to these analysis histories, as well as the one-click workflow itself, are shown in **Table 3** below. Analysis of the published data generated six verified neoantigen candidate peptides predicted from String-Tie identified non-reference sequences (Item 153 in the shared analysis history), but none from CDS sequences (SAVs or InDels).

**Table 3.**
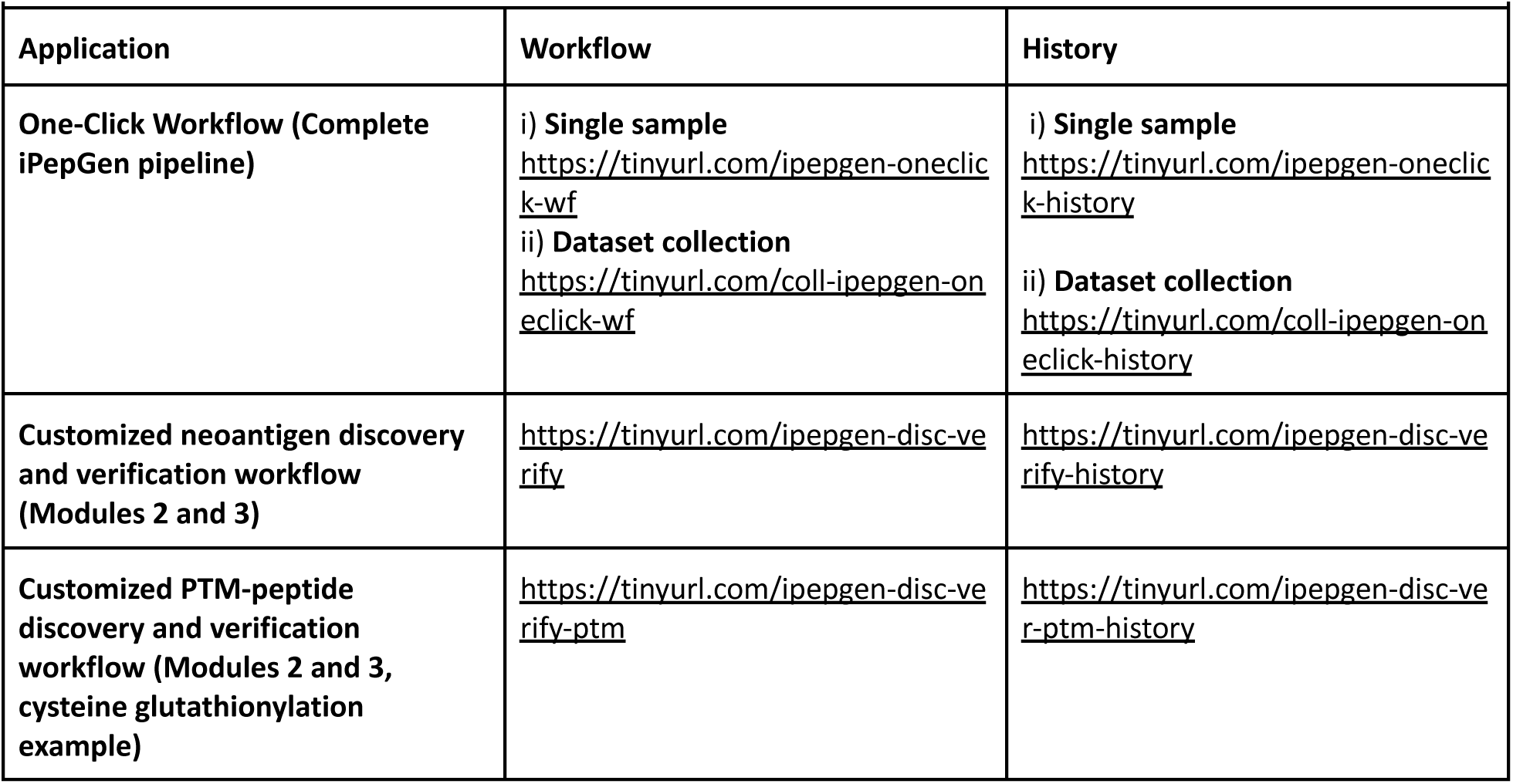
Links to workflows and demonstration analysis histories for the One-Click Workflow and Customized Workflows.

#### Flexibility of iPepGen: building customized workflows for neoantigen identification and verification, and PTM analysis

Galaxy provides the unique flexibility to integrate selected modules of iPepGen into customized workflows tailored to specific analysis applications. One use-case requiring iPepGen customization would be scenarios where a customized protein FASTA database (containing potential non-reference neoantigen sequences) has been generated externally, and PSM discovery and verification from MS/MS data is required. For such a situation, we built a customized workflow that integrates Modules 2 and 3 for neoantigen discovery and verification, respectively.

To demonstrate the effectiveness of this customized iPepGen workflow, we selected data from a published study by the Bassani-Sternberg group focused on neoantigen discovery in a melanoma cell line [90]. We selected four biological replicate immunopeptidomics raw MS/MS datasets generated from untreated control melanoma cells and four raw MS/MS files from biological replicates of cells treated with 5-aza-2′-deoxycytidine (DAC), which promotes processing and presentation of peptide antigens on the HLA class I complex. These raw MS/MS datasets were generated using optimized immunopeptide enrichment and MS instrumental analysis methods, such that they represented extremely data-rich MS/MS files for demonstrating the effectiveness of this customized workflow. A customized protein sequence database of about 900,000 sequences made available in the published data, was used. It contained predicted, novel proteoform sequences from long-noncoding RNA, transposable elements, and other non-reference sequences, including SAVs and other CDS mutations, annotated from RNA-Seq data on these cell lines. Analysis by our customized pipeline identified 22,386 distinct peptides using FragPipe across these eight MS/MS files in Module 2, complete with LFQ abundance levels (item 49 in the linked analysis history in **Table 3**). This deep dataset also generated several hundred potential neoantigen candidates, which were advanced to the verification Module using PepQuery2 and BLASTP. 93 distinct neoantigen candidate peptides were verified (item 85 in the linked analysis history in **Table 3).** We were able to quantify neoantigen candidates and their response to DAC treatment, as well as known TAAs derived from reference human protein sequences. **Supplemental Figure 7** shows example results for a verified neoantigen peptide and a known melanoma TAA, which showed an expected increased abundance with DAC treatment.

We also used this customized workflow to demonstrate the ability of the iPepGen pipeline to identify HLA-bound peptides carrying PTMs that may regulate peptide processing for presentation to the immune system. For this demonstration, we focused on cysteine glutathionylation, known to regulate HLA presentation [91]. We used six MS/MS datafiles from the same published study analyzing melanoma cells [90], and added cysteine glutathionylation as a variable modification for the FragPipe analysis. This resulted in the identification of 105 peptides containing glutathionylated cysteine (item 41 in the linked history in **Table 3**), complete with LFQ abundance measures using IonQuant in FragPipe. These candidates were then passed on to PepQuery2 for verification. Here, PepQuery2 was operated to evaluate “known” sequences in the human reference proteome, including those with a variable modification to cysteine. PepQuery verified 28 distinct cysteine glutathionylated (item 64, the linked analysis history in **Table 3**).

## Discussion

iPepGen’s design addresses key challenges unique to immunopeptidomics, including the use of personalized protein databases derived from NGS data, peptide identification using no enzyme specificity, verification of non-reference peptide sequences, and computational prioritization of candidate neoantigens. Several other informatics platforms aimed at complete immunopepetidogenomic analysis have also been described [42,47,48], notably the NeoDisc [51] pipeline, which offers a sophisticated set of tools including a deep learning algorithm to prioritize candidate neoantigens with the highest potential for value in immunotherapy. iPepGen offers similar capabilities to these available platforms, but also several distinguishing features:

● iPepGen implementation in the Galaxy ecosystem, featuring accessibility on publicly available, scalable cloud-computing infrastructure, enables usage by those who lack software developer expertise, and/or access to HPC computing environments required for compute-intensive applications inherent to immunopeptidogenomics. This contrasts with platforms such as NeoDisc, which, despite offering very powerful capabilities including the use of FragPipe for PSM discovery, require advanced technical know-how and access to local HPC infrastructure for implementation and maintenance of its operation, limiting its use by those lacking this expertise and computer hardware.
● The availability of online training material through the GTN empowers adoption by users with limited computational expertise, providing a step-by-step guide to learn each module of the iPepGen pipeline in a hands-on manner, using supplied demonstration data. Once trained, these users can advance to using the One-Click workflow for analysis of their own datasets.
● Trained users can take advantage of the transparent and flexible features of the Galaxy analysis environment to customize their own analyses, using the described iPepGen modules as a foundation. Users can take advantage of the rich library of training material offered by the GTN [56,57] to integrate new software tools (e.g., additional tools to predict novel proteoforms from genomic data) or build customized workflows tailored to their analysis needs. Adaptability to new software integration is a hallmark of the Galaxy ecosystem, which also offers a thriving community of developers and users, ensuring sustained maintenance and operation [53]. Other platforms, such as NeoDisc, do not offer such flexibility and require advanced software development expertise to customize the pipeline and maintain its operation.

In **Module 1**, we have developed several workflows to assemble and annotate a diverse profile of non-reference genomic and transcriptomic sequences that may give rise to novel proteoforms, which, in turn, are potentially processed into neoantigen peptides. We have developed these workflows to build protein databases personalized to the user’s sample(s) with a broad profile of potential neoantigen sequences, which are confirmed via integration with MS/MS data in Module 2. This module is flexible to customization with other approaches to identifying specific types of non-reference genomic and/or transcriptomic sequences from NGS data, including single-cell, long-read sequencing, or other “third-generation sequencing” methods [92]. Galaxy has a well-established history as an environment amenable to the integration of emerging software for genomic and transcriptomic sequencing analysis [93]. iPepGen offers compatibility with a wide array of analysis tools in this area, providing a means to build protein sequence databases containing potential neoantigen sequences for interrogation with immunopeptidomic MS/MS data.

The use of FragPipe in **Module 2**, and specifically the MSFragger algorithm, provides the backbone for matching immunopeptidomic MS/MS data to the customized sequence databases from Module 1. MSFragger’s ability to efficiently handle “no enzyme” search settings is critical in this context, as MHC-bound peptides are often produced by a variety of proteases, unlike typical proteomic samples that are digested with trypsin. The expanded search space inherent to immunopeptidomic data substantially increases memory and processing requirements, especially when searching against a custom database containing over 2 million protein sequences. To accommodate this, we enabled MSFragger’s “split database” functionality, dividing the database into 200 sections. Deployment of iPepGen in a scalable, cloud-based Galaxy instance is also key. The European Galaxy instance housing our workflows allocates up to 250 GB of active memory, which proved sufficient for executing MSFragger under these conditions.

We also evaluated the MSBooster tool [66] in FragPipe, which utilizes machine-learning-based scoring to predict fragment ion intensities and retention times. After this prediction, potential PSMs are re-evaluated, which results in the rescue of many peptide identifications and enhances sensitivity. In our analysis of representative datasets (**Figure 3B** and **Supplemental Figure 2**), incorporating MSBooster significantly increased identification of peptides, both reference and non-reference neoantigen candidates, without extending runtime or memory consumption. iPepGen also takes advantage of FragPipe’s IonQuant module [46] for LFQ by aligning peptides across LC-MS/MS datasets and calculating abundances based on area-under-the-curve measurements and match-between-run functionality. This feature is enabled in our shared workflows and allows researchers to incorporate abundance measurements of identified neoantigen peptides and compare their response across different experimental conditions.

In **Module 3**, PepQuery2 offers a peptide-centric means to verify candidate neoantigen sequences through a secondary analysis of peptides discovered in Module 2. Unlike conventional sequence database search tools, PepQuery2 focuses exclusively on user-defined candidate neoantigen peptide sequences and evaluates their match quality to potential MS/MS spectra in the dataset against human reference peptides, while also accounting for common PTMs to these peptides. Our use of PepQuery2 is another distinguishing feature of iPepGen, as other pipelines rely on traditional sequence database searching to identify neoantigen candidates and lack such a stringent verification step as that offered by PepQuery2. We did observe that using non-specific enzyme digestion increases memory demands for PepQuery2, such that limiting the amino acid length of reference peptides for consideration to a narrow window around the input neoantigen candidates was necessary for successful operation (see Methods for more details).

**Module 4** provides an initial means to classify non-reference peptides based on the aberrant genomic or transcriptomic sequences coding for the identified sequence. The simple tabular output also provides links to be imported into the popular IGV or UCSC genome browser to visualize the genomic coordinates coding for the peptide, along with any supporting transcript sequences responsible for the translation product. As we note in the Results section describing this module, and **Supplemental Figure 5**, the complex nature of some aberrant genomic and/or transcriptomic sequences coding neoantigen peptides may require deeper manual investigation to understand the source of some peptide sequences. The ability to visualize the sequences coding for candidate neoantigen peptides in IGV or UCSC offers users guidance in characterizing these complex scenarios.

The HLA genotyping tools in **Module 5**, OptiType and Seq2HLA, provide complementary strengths for sequence analysis of HLA alleles from NGS data. OptiType is optimized for Class I alleles, while Seq2HLA genotypes both Class I and Class II alleles. Use of these two tools offers more comprehensive genotyping of HLA alleles. While Module 5 provides basic prioritization functionality using prediction of binding affinity to the genotyped alleles as a factor in immunogenicity, more advanced machine-learning-based tools are increasingly available to predict immunogenicity from multi-omics datasets [24,94–96]. With this in mind, we have implemented the pVACbind [87] software in Galaxy, part of the pVACtools software suite [24], compatible with the outputs generated by our pipeline. pVACbind provides complementary tools for predicting HLA binding, as well as two deep learning algorithms for predicting immunogenicity [29,30].

In addition to the availability of workflows and accompanying training material making up each module of iPepGen, we have also developed several customized versions of the pipeline. These demonstrate the flexibility offered by Galaxy implementation. One of these is the “One-click” workflow, which offers a more streamlined option for advanced users to upload appropriate raw data and generate results in a single click. We have demonstrated this workflow’s effectiveness on data available from a published immunopeptidogenomics study [89]. It should offer an easy-to-use workflow for users who have gained training via our GTN materials and are ready to analyze their own data. We also developed a customized workflow that integrates Modules 2 and 3, for use-cases where a FASTA-formatted protein sequence database containing potential non-reference neoantigen sequences is already available, along with immunopeptidomics MS/MS data. We demonstrate the effectiveness of this workflow on deep immunopeptidomics data from a published study [90], showing the ability of iPepGen to efficiently handle larger volumes of data. Finally, we use this customized workflow and this same dataset to demonstrate the ability to identify and verify PTMs on HLA-bound peptides, using cysteine glutathionylation as an example. Collectively, these extensions of iPepGen provide a demonstration of the unique flexibility of the pipeline to adjust to the experimental needs of users.

Although iPepGen offers a rich set of functionalities required for immunopeptidogenomic data analysis, some additional features will be the focus of future work. We have focused our demonstration of its effectiveness on HLA class I immunopeptidomics data. However, the pipeline can be easily adapted to class II studies. Module 2 already generates PSMs for a broad range of peptide lengths, including the most common lengths (12-20 amino acids) for class II-bound peptides [97]. This length range can be modified in FragPipe as necessary if other peptide lengths are anticipated. In Module 3, the length range for PepQuery2 can be easily adjusted to verify peptides within this length range. Seq2HLA offers the ability to genotype HLA class II alleles, and IEDB tools in Galaxy can predict the binding of peptides to these alleles. Moreover, pVACbind provides several additional tools for class II peptide binding prediction, and includes the DeepImmuno tool [29] that predicts immunogenicity for class II peptide-allele pairs.

The current workflow focuses on the analysis of data-dependent acquisition (DDA) peptide MS/MS data. With the advent of new, faster scanning instruments, data-independent acquisition (DIA) data is increasingly used for immunopeptidomics [98,99]. DIA offers potential for deeper detection of peptides in complex mixtures and more accurate and precise quantification [100]. The FragPipe suite offers tools for DIA analysis [101,102], which are readily available for analysis of these data types in our pipeline. However, utilizing a large, customized sequence database and verifying neoantigen candidates discovered from DIA data offers challenges that will need to be addressed. Lastly, de novo sequencing from MS/MS data could offer identification of neoantigen peptide candidates that are not predicted from genomic or transcriptomic data [103], and could be explored for added value.

## Conclusion

In summary, the iPepGen pipeline offers several distinguishing qualities from other available informatics solutions for immunopeptidogenomic informatics:

● Immediate access to analysis modules on a scalable cloud-based infrastructure, with the option to run individual modules or together as a complete pipeline for the discovery, verification, and prioritization of neoantigen peptides for further experimental evaluation.
● Step-by-step, interactive training material for each module is encapsulated in a customized learning pathway [58], seamlessly linked to accessible workflows, empowering new users with limited bioinformatic expertise or access to advanced computational resources to learn and use the pipeline for their studies.
● Workflows composed of tools selected to meet key challenges and common bottlenecks of neoantigen discovery and verification, offering efficient and sensitive identification of peptides processed by non-specific proteases and rigorous verification of candidate neoantigen peptides using two algorithms (FragPipe and PepQuery2).
● Extensible and customizable implementation, open to new software tools as they emerge, and development of customized workflows integrating selected modules of the pipeline to meet user analysis requirements.
● Promotion of reproducibility and transparency via sharing of customized workflows and completed analysis histories for use by others and/or as an accessible record to accompany method descriptions in peer-reviewed publications.

Taken together, these features of iPepGen make it a unique and timely resource for the growing community of researchers conducting immunopeptidogenomics analyses. It offers an important contribution to the immuno-oncology research community seeking to develop improved therapies for treating and preventing a wide range of cancers.

## Methods

### Aim, Design, and Setting of the Study

This study aimed to develop and validate iPepGen—a modular, end-to-end, Galaxy-based immunopeptidogenomic pipeline designed to enable accessible and reproducible identification, verification, and prioritization of neoantigen peptides from cancer multi-omics data. The design integrates genomic, transcriptomic, and mass spectrometry (MS)-based immunopeptidomics data. Development of each module and associated workflows, along with demonstration data analysis, was conducted on the European Galaxy server [104].

### Description of Materials and Data

For data demonstrating the effectiveness of iPepGen analysis modules, we used the STS-26T malignant peripheral nerve sheath tumor (MPNST) cell line. Total RNA was extracted and sequenced on an Illumina platform, yielding 100 base-pair paired-end reads (∼80 million reads/sample). Cells were treated with gentamicin (50 µg/mL) to promote ribosomal read-through and enhance expression of non-reference proteoforms [105]. MHC class I complexes were immunoprecipitated using the W6/32 monoclonal antibody, and bound peptides were analyzed on an Orbitrap Eclipse mass spectrometer using data-dependent acquisition (DDA) mode [106,107]. Using RNA-Seq and immunopeptidomics data from an MPNST cell line as a representative of user data inputs and analysis requirements, we describe the functionality and show representative results demonstrating the effectiveness of the iPepGen pipeline. **Table 1** provides links to the raw input data generated from the MPNST cells, as well as completed analysis histories.

### iPepGen Pipeline Modules

The links in Table 1 provide access to the Galaxy software tools composing workflows of each iPepGen module, along with validated and optimized settings for their operation. Here, we describe briefly the core tools used in each workflow and the settings for specific tools necessary for effective operation and performance.

#### Module 1: Prediction of Non-reference Neoantigen Peptides

Paired-end RNA-Seq reads are aligned to the GRCh38 reference genome using HISAT2 [61]. In one workflow, gene fusion events are identified using Arriba [59], which utilizes spliced alignments from STAR [60] to detect chimeric reads. For broader non-canonical transcript detection, including single-nucleotide variants (SAVs), indels, and aberrant transcripts, a workflow using FreeBayes [108] was employed for variant calling coupled with CustomProDB [62], generating a protein database of SAV and short InDel sequences. A parallel workflow uses StringTie [63] and GffCompare [64] for transcript assembly and annotation of non-reference sequences, followed by the use of the Galaxy tool “Translate BED transcripts” and “bed to protein map” to generate a customized FASTA database. For both the CustomProDB and StringTie-based workflows, permissive thresholds for Reads Per Kilobase per Million mapped reads (RPKM) were used (Value of 1 for CustomProDB and 2 for StringTie) to retain low-abundance transcripts that may encode tumor-specific proteoforms. Customized protein databases with predicted non-reference sequences are merged along with the human reference sequences from UniProt [109] and common contaminant proteins using the Galaxy tool “fasta_merge_files_and_filter_unique_sequences”, generating a comprehensive, annotated FASTA-formatted database of over 2.9M sequences for downstream analysis.

This module requires the installation and configuration of HISAT2, STAR, FreeBayes, StringTie, GffCompare, Arriba, and CustomProDB within Galaxy. Of these, FreeBayes is the most computationally intensive, requiring approximately 10 CPUs and at least 17 GB of RAM per sample. A single RNA-Seq dataset may take up to 30 hours to process. Other tools in the module typically operate efficiently with 4–8 CPUs and 8–16 GB RAM, completing in less than 4–6 hours.

#### Module 2: Discovery of Candidate Neoantigen Peptides

In module 2, raw LC-MS/MS files (for demonstration data, .raw formatted files from the ThermoFisher Orbitrap instrument) can be converted to mzML format using msconvert [110] or directly analyzed in native format by the FragPipe software suite. For this module, FragPipe (v23.0) uses these software components: MSFragger [70] for sequence database searching under no-enzyme settings (allowing identification of peptides 8–20 amino acids long), MSBooster [66] for deep-learning-based prediction of fragmentation and retention time patterns, Percolator [67] for machine-learning-based PSM validation, and Philosopher [68] for false discovery rate (FDR) estimation using a target-decoy approach. For demonstration data from MPNST cells, MS/MS spectra generated on the Orbitrap LC-MS/MS system were matched to the merged protein FASTA database generated in Module 1, generating peptide spectrum matches (PSMs). For the demonstration data, only oxidation on methionine was included as a variable PTM. Non-reference peptides were first automatically selected based on lack of direct sequence match to a known reference protein provided in the tabular FragPipe peptide report, and then extracted with annotation using Galaxy’s Query Tabular tool [69], based on header annotations in the FASTA file indicating their origin from the workflows in Module 1 (e.g., fusion, SAV/InDel, or novel transcript).

This module requires the Galaxy-deployed FragPipe suite, msconvert, and Query Tabular. PSM generation using non-specific enzyme settings against a large customized sequence database of several million entries is memory intensive. To address this challenge, it is important to configure FragPipe to use a database split of at least 200 to lessen the memory burden. Users could also disable the “Mass calibration and optimization” setting as well to save on memory load. Additionally, the Galaxy FragPipe tool was configured with an allocation of 16 CPUs and 250 GB RAM. Using these settings, a typical DDA file (∼1 GB) was processed within 6 to 8 hours on the European Galaxy server.

#### Module 3: Verification of Neoantigen Peptide-Spectrum Matches

This module implements a second level of PSM validation using PepQuery2 [71]. Neoantigen candidate peptides identified in Module 2 input as a tabular list of sequences are re-evaluated against the raw MS/MS spectral data, converted to MGF format via msconvert, and the UniProt reference proteome. PepQuery2 conducts peptide-centric analysis by considering reference proteome sequences of close length and mass to the candidate neoantigen sequence, along with common PTMs (e.g., phosphorylation, deamidation, acetylation, etc.), as potentially competing matches to MS/MS spectra that led to the identification of the non-reference sequence. PepQuery also generates thousands of randomized sequences of similar amino acid length and mass and evaluates potential matches to the MS/MS spectra under consideration. Through these evaluations, a hyperscore is assigned for sequences matching the MS/MS spectrum, and statistical modeling to assign p-values to the best sequence match to the spectrum is conducted. Verified neoantigen peptides are those that remain the top match to the MS/MS spectrum (as opposed to a reference sequence or a randomized sequence) and are assigned a p-value indicating strong statistical significance. PepQuery2 assigns a Yes or No Confidence to each input peptide based on these assigned values. In our workflow, neoantigen peptides deemed confident are further evaluated using BLAST-P against the NCBI reference database to confirm that these are truly novel sequences, with no exact matching sequences found in the reference proteome. We note that the non-redundant NCBI reference database is continuously updated, so the output novel sequences from this module may change over time with these updates.

To ensure the performance of PepQuery2, when considering peptides from non-specific enzyme processing, using narrow amino acid length windows for reference sequences to consider is critical. For example, for neoantigen candidate peptide sequences nine amino acids in length, a narrow window (e.g., 9-11 amino acids) is suggested to minimize the number of reference peptides that need to be generated initially by the software, controlling the active memory required. These settings provide the algorithm with thousands of reference peptides to consider as competing for matching to MS/MS. Only peptides falling into the amino acid length range selected by the user can be confidently verified, so any input peptides of other lengths should be ignored. Using these optimized conditions, PepQuery2 completed the evaluation of 123 candidate sequences in under 20 minutes, using ≤8 GB RAM and 2 CPUs. BLAST-P queries using the Galaxy tool were also completed in under 30 minutes using similar compute allocations, making this module both computationally tractable and scalable.

#### Module 4: Visualization and Classification of Neoantigen Origins

This module utilizes the PepPointer tool [74] to map non-reference neoantigen peptides to their genomic coordinates using BED and GTF annotations assigned to the sequences. It also classifies their origin as being from coding DNA sequence (CDS), untranslated regions (UTR), intron-retained, or other intergenic sequences. The tabular output provides links with necessary information to be input into the IGV [75] or the UCSC Genome Browser [76], enabling researchers to inspect peptide-transcript-genome alignments visually. Galaxy’s Query Tabular tool was used to format, annotate, and extract relevant features from intermediate outputs and generate tabular results with formats compatible with PepPointer. The tools in this module are not compute-intensive, typically completing in <15 minutes using 1–2 CPUs and ≤4 GB RAM, even when analyzing multiple peptide entries.

#### Module 5: Prioritization of Neoantigens via HLA Binding Prediction

HLA genotyping is performed using two complementary tools: OptiType [83], optimized for class I alleles, and Seq2HLA [84], which also covers class II loci. Paired RNA-Seq fastq files serve as input, and genotype outputs are merged using AWK in Galaxy and formatted via Query Tabular into tabular formats compatible with the IEDB MHC Binding Prediction tool [85]. The NetMHCpan_EL algorithm, trained on empirical data for eluted peptides from immunopeptidomics studies, predicts binding affinity of verified neoantigen peptides (which are provided as input) to specific HLA alleles. Peptides are classified as strong (binding affinity percentile rank ≤ 0.5%) or weak binders (percentile rank 0.5–2%) to specific HLA sequences. Peptides of 9 or more amino acids are processed into shorter sub-sequences by the IEDB tools, and their binding affinity to HLA alleles is also assessed. However, only the peptides verified by PepQuery in Module 3 have evidence supporting their presence in the sample(s) and are outputted in the final report on peptide binders.

The outputs of the pipeline (FASTA-formatted neoantigen peptide candidates and a tabular list of HLA genotypes) can also be used as inputs to the pVACbind software in Galaxy. The user can select from a portfolio of tools in pVACbind for predicting HLA allele binding, and also immunogenicity using deep learning algorithms. The output is a tabular file, which shows all predicted values from each selected algorithm. Users can consult pVACbind documentation to assist in the interpretation of these results [87] and **Supplemental Figure 6(a,b,c)** for guidance on its operation in Galaxy.

This module requires Galaxy-deployed OptiType, Seq2HLA, the IEDB binding prediction software, PepQuery2 (optional), and native Galaxy text processing tools (AWK, Query Tabular). OptiType required 8 CPUs and 32 GB RAM, with runtimes of ∼32 hours, while Seq2HLA required 4 CPUs and 16 GB RAM, completing in ∼24 hours. The IEDB predictor operated efficiently with 2–4 CPUs and ≤8 GB RAM, processing ∼100 peptides in ∼30 minutes.

## Abbreviations

Browser Extensible Data (BED) formatted file

common Repository of Adventitious Proteins (cRAP)

coding DNA sequence (CDS)

5-aza-2’-deoxycytidine (DAC)

data-dependent acquisition (DDA)

data-independent acquisition (DIA) eluted ligand (EL)

false-discovery rate (FDR)

Galaxy Training Network (GTN)

Gene Transfer File (GTF)

high-performance computing (HPC)

human leukocyte antigen (HLA)

immunopeptidogenomics (iPepGen)

Insertions and Deletion (InDel)

Integrated Genome Viewer (IGV)

label-free quantification (LFQ)

liquid chromatography (LC)

major histocompatibility complex (MHC)

malignant peripheral nerve sheath tumor (MPNST)

Mascot generic format (.mgf)

mass spectrometry (MS)

National Library of Medicine-National Center for Biotechnology Information (NLM-NCBI)

Next-generation sequencing (NGS)

personalized Variant Antigens in Cancer tool suite (pVACtools)

post-translational modification (PTM)

reads per kilobase per million mapped reads (RPKM)

single amino acid variants (SAVs)

tandem mass spectrometry (MS/MS)

tumor-associated antigen (TAA)

Tumor-specific antigen (TSA)

untranslated region (UTR)

## Declarations

### Ethics approval and consent to participate

Not applicable

### Consent for publication

Not applicable

### Availability of data and materials

All the software tools composing the iPepGen pipeline and its workflow modules are available through the Galaxy Tool Shed [111]. The Tool Shed page for each software tool also provides links to GitHub repositories with source code and documentation, along with license information. Most software used in the iPepGen modules is developed using permissive licensing (e.g., GNU GPL, MIT). Some of the tools contained within the FragPipe software platform (MSFragger and the associated IonQuant software are developed under an academic license and are available freely for academic research, non-commercial, or educational purposes. Other software contained in FragPipe is developed under permissive licensing [112]. The Galaxy bioinformatics platform is freely available and open-source, released under the MIT license [113]. All data, including representative input files, intermediate processed results, and final outputs, are available on the publicly accessible European Galaxy server, as described in **Table 1**. As well as users can also download the data and resultant files from the MassIVE repository MSV000100019 or from PRIDE (PXD071206).

### Authors’ contributions

S.M. conceptualized the study, curated data, performed formal analysis, developed software, created visualizations, and drafted and revised the manuscript. R.W. contributed to data curation, software development, and manuscript review. K.T.D. participated in data curation, formal analysis, software development, and manuscript editing. J.E.J. contributed to conceptualization, data curation, and software implementation. F.Y. provided data resources, contributed to software development, and participated in manuscript review. T.J. supported data acquisition and manuscript review. K.R. contributed resources, software support, and manuscript editing. S.C. provided resources and reviewed the manuscript. F.P. assisted in data curation, resource provision, and manuscript editing. A.I.N. contributed software, supervision, funding acquisition, and manuscript review. D.A.L. led conceptualization, resource acquisition, supervision, funding, and manuscript review. P.D.J. contributed to conceptualization, software development, supervision, funding acquisition, and manuscript review. T.J.G. led conceptualization, software development, supervision, and funding acquisition, and contributed to drafting and revising the manuscript. All authors read and approved the final manuscript.

## Acknowledgements

We thank the Center for Metabolomics and Proteomics at the University of Minnesota for providing services to generate MS-based immunopeptidomics data used for developing, testing, and producing training resources for the iPepGen pipeline as described in the manuscript. We also thank Dr. Bing Zhang (Baylor College of Medicine, Houston, TX) for advice on the use of PepQuery2 within our pipeline, and Dr. Malachi Griffith (Washington University School of Medicine, St. Louis, MO) for assistance in the use of pVACbind.

## References

1. Wang Y, Shi T, Song X, Liu B, Wei J. Gene fusion neoantigens: Emerging targets for cancer immunotherapy. Cancer Lett. 2021;506:45–54. 10.1016/j.canlet.2021.02.023

2. Laumont CM, Vincent K, Hesnard L, Audemard É, Bonneil É, Laverdure J-P, et al. Noncoding regions are the main source of targetable tumor-specific antigens. Sci Transl Med. 2018;10:eaau5516. 10.1126/scitranslmed.aau5516

3. Yi X, Liao Y, Wen B, Li K, Dou Y, Savage SR, et al. caAtlas: An immunopeptidome atlas of human cancer. iScience. 2021;24:103107. 10.1016/j.isci.2021.103107

4. Wang E, Aifantis I. RNA Splicing and Cancer. Trends Cancer. 2020;6:631–44. 10.1016/j.trecan.2020.04.011

5. Smith LM, Kelleher NL. Proteoform: a single term describing protein complexity. Nat Methods. 2013;10:186–7. 10.1038/nmeth.2369

6. Capietto A-H, Hoshyar R, Delamarre L. Sources of Cancer Neoantigens beyond Single-Nucleotide Variants. Int J Mol Sci. 2022;23:10131. 10.3390/ijms231710131

7. Xie N, Shen G, Gao W, Huang Z, Huang C, Fu L. Neoantigens: promising targets for cancer therapy. Signal Transduct Target Ther. 2023;8:9. 10.1038/s41392-022-01270-x

8. Ahn R, Cui Y, White FM. Antigen discovery for the development of cancer immunotherapy. Semin Immunol. 2023;66:101733. 10.1016/j.smim.2023.101733

9. Minati R, Perreault C, Thibault P. A Roadmap Toward the Definition of Actionable Tumor-Specific Antigens. Front Immunol. 2020;11:583287. 10.3389/fimmu.2020.583287

10. Archbold JK, Macdonald WA, Gras S, Ely LK, Miles JJ, Bell MJ, et al. Natural micropolymorphism in human leukocyte antigens provides a basis for genetic control of antigen recognition. J Exp Med. 2009;206:209–19. 10.1084/jem.20082136

11. Kaufman JF, Auffray C, Korman AJ, Shackelford DA, Strominger J. The class II molecules of the human and murine major histocompatibility complex. Cell. 1984;36:1–13. 10.1016/0092-8674(84)90068-0

12. Chen DS, Mellman I. Elements of cancer immunity and the cancer-immune set point. Nature. 2017;541:321–30. 10.1038/nature21349

13. Kim SK, Cho SW. The Evasion Mechanisms of Cancer Immunity and Drug Intervention in the Tumor Microenvironment. Front Pharmacol. 2022;13:868695. 10.3389/fphar.2022.868695

14. Blass E, Ott PA. Advances in the development of personalized neoantigen-based therapeutic cancer vaccines. Nat Rev Clin Oncol. 2021;18:215–29. 10.1038/s41571-020-00460-2

15. Chen F, Zou Z, Du J, Su S, Shao J, Meng F, et al. Neoantigen identification strategies enable personalized immunotherapy in refractory solid tumors. J Clin Invest. 129:2056–70. 10.1172/JCI99538

16. Braun DA, Moranzoni G, Chea V, McGregor BA, Blass E, Tu CR, et al. A neoantigen vaccine generates antitumour immunity in renal cell carcinoma. Nature. 2025;639:474–82. 10.1038/s41586-024-08507-5

17. Zhang X, Goedegebuure SP, Chen MY, Mishra R, Zhang F, Yu YY, et al. Neoantigen DNA vaccines are safe, feasible, and induce neoantigen-specific immune responses in triple-negative breast cancer patients. Genome Med. 2024;16:131. 10.1186/s13073-024-01388-3

18. Redwood AJ, Dick IM, Creaney J, Robinson BWS. What’s next in cancer immunotherapy? - The promise and challenges of neoantigen vaccination. Oncoimmunology. 11:2038403. 10.1080/2162402X.2022.2038403

19. Andersen MH. Tumor microenvironment antigens. Semin Immunopathol. 2023;45:253–64. 10.1007/s00281-022-00966-0

20. Bais P, Namburi S, Gatti DM, Zhang X, Chuang JH. CloudNeo: a cloud pipeline for identifying patient-specific tumor neoantigens. Bioinformatics. 2017;33:3110–2. 10.1093/bioinformatics/btx375

21. Chai S, Smith CC, Kochar TK, Hunsucker SA, Beck W, Olsen KS, et al. NeoSplice: a bioinformatics method for prediction of splice variant neoantigens. Bioinform Adv. 2022;2:vbac032. 10.1093/bioadv/vbac032

22. Hundal J, Carreno BM, Petti AA, Linette GP, Griffith OL, Mardis ER, et al. pVAC-Seq: A genome-guided in silico approach to identifying tumor neoantigens. Genome Med. 2016;8:11. 10.1186/s13073-016-0264-5

23. Vensko SP, Olsen K, Bortone D, Smith CC, Chai S, Beckabir W, et al. LENS: Landscape of Effective Neoantigens Software. Bioinformatics. 2023;39:btad322. 10.1093/bioinformatics/btad322

24. Hundal J, Kiwala S, McMichael J, Miller CA, Xia H, Wollam AT, et al. pVACtools: A Computational Toolkit to Identify and Visualize Cancer Neoantigens. Cancer Immunol Res. 2020;8:409–20. 10.1158/2326-6066.CIR-19-0401

25. Dhanda SK, Mahajan S, Paul S, Yan Z, Kim H, Jespersen MC, et al. IEDB-AR: immune epitope database—analysis resource in 2019. Nucleic Acids Res. 2019;47:W502–6. 10.1093/nar/gkz452

26. Kalemati M, Darvishi S, Koohi S. CapsNet-MHC predicts peptide-MHC class I binding based on capsule neural networks. Commun Biol. 2023;6:492. 10.1038/s42003-023-04867-2

27. Koşaloğlu-Yalçın Z, Blazeska N, Vita R, Carter H, Nielsen M, Schoenberger S, et al. The Cancer Epitope Database and Analysis Resource (CEDAR). Nucleic Acids Res. 2022;51:D845–52. 10.1093/nar/gkac902

28. Calis JJA, Maybeno M, Greenbaum JA, Weiskopf D, De Silva AD, Sette A, et al. Properties of MHC Class I Presented Peptides That Enhance Immunogenicity. PLoS Comput Biol. 2013;9:e1003266. 10.1371/journal.pcbi.1003266

29. Li G, Iyer B, Prasath VBS, Ni Y, Salomonis N. DeepImmuno: deep learning-empowered prediction and generation of immunogenic peptides for T-cell immunity. Brief Bioinform. 2021;22:bbab160. 10.1093/bib/bbab160

30. Albert BA, Yang Y, Shao XM, Singh D, Smit KN, Anagnostou V, et al. Deep neural networks predict class I major histocompatibility complex epitope presentation and transfer learn neoepitope immunogenicity. Nat Mach Intell. 2023;5:861–72. 10.1038/s42256-023-00694-6

31. Elfatimi E, Lekbach Y, Prakash S, BenMohamed L. Artificial intelligence and machine learning in the development of vaccines and immunotherapeutics-yesterday, today, and tomorrow. Front Artif Intell. 2025;8:1620572. 10.3389/frai.2025.1620572

32. Becker JP, Riemer AB. The Importance of Being Presented: Target Validation by Immunopeptidomics for Epitope-Specific Immunotherapies. Front Immunol. 2022;13:883989. 10.3389/fimmu.2022.883989

33. Kacen A, Javitt A, Kramer MP, Morgenstern D, Tsaban T, Shmueli MD, et al. Post-translational modifications reshape the antigenic landscape of the MHC I immunopeptidome in tumors. Nat Biotechnol. 2023;41:239–51. 10.1038/s41587-022-01464-2

34. Zeng H, Gifford DK. Quantification of uncertainty in peptide-MHC binding prediction improves high-affinity peptide selection for therapeutic design. Cell Syst. 2019;9:159–166.e3. 10.1016/j.cels.2019.05.004

35. Zhao W, Sher X. Systematically benchmarking peptide-MHC binding predictors: From synthetic to naturally processed epitopes. PLoS Comput Biol. 2018;14:e1006457. 10.1371/journal.pcbi.1006457

36. Feola S, Chiaro J, Martins B, Russo S, Fusciello M, Ylösmäki E, et al. A novel immunopeptidomic-based pipeline for the generation of personalized oncolytic cancer vaccines. eLife. 11:e71156. 10.7554/eLife.71156

37. Gfeller D, Liu Y, Racle J. Contemplating immunopeptidomes to better predict them. Semin Immunol. 2023;66:101708. 10.1016/j.smim.2022.101708

38. Shapiro IE, Bassani-Sternberg M. The impact of immunopeptidomics: From basic research to clinical implementation. Semin Immunol. 2023;66:101727. 10.1016/j.smim.2023.101727

39. Terai YL, Huang C, Wang B, Kang X, Han J, Douglass J, et al. Valid-NEO: A Multi-Omics Platform for Neoantigen Detection and Quantification from Limited Clinical Samples. Cancers (Basel). 2022;14:1243. 10.3390/cancers14051243

40. Hunt DF, Henderson RA, Shabanowitz J, Sakaguchi K, Michel H, Sevilir N, et al. Pillars article: Characterization of peptides bound to the class I MHC molecule HLA-A2.1 by mass spectrometry. Science 1992. 255: 1261-1263. J Immunol. 2007;179:2669–71.

41. Rivero-Hinojosa S, Grant M, Panigrahi A, Zhang H, Caisova V, Bollard CM, et al. Proteogenomic discovery of neoantigens facilitates personalized multi-antigen targeted T cell immunotherapy for brain tumors. Nat Commun. 2021;12:6689. 10.1038/s41467-021-26936-y

42. Scull KE, Pandey K, Ramarathinam SH, Purcell AW. Immunopeptidogenomics: Harnessing RNA-Seq to Illuminate the Dark Immunopeptidome. Mol Cell Proteomics. 2021;20:100143. 10.1016/j.mcpro.2021.100143

43. Sandalova T, Sala BM, Achour A. Structural aspects of chemical modifications in the MHC-restricted immunopeptidome; Implications for immune recognition. Front Chem. 2022;10:861609. 10.3389/fchem.2022.861609

44. Capturing the diversity of protein modifications on presented tumor antigens. Nat Biotechnol. 2023;41:195–6. 10.1038/s41587-022-01465-1

45. Mehta S, Easterly CW, Sajulga R, Millikin RJ, Argentini A, Eguinoa I, et al. Precursor Intensity-Based Label-Free Quantification Software Tools for Proteomic and Multi-Omic Analysis within the Galaxy Platform. Proteomes. 2020;8:15. 10.3390/proteomes8030015

46. Yu F, Haynes SE, Nesvizhskii AI. IonQuant Enables Accurate and Sensitive Label-Free Quantification With FDR-Controlled Match-Between-Runs. Mol Cell Proteomics. 2021;20:100077. 10.1016/j.mcpro.2021.100077

47. Li Y, Wang G, Tan X, Ouyang J, Zhang M, Song X, et al. ProGeo-neo: a customized proteogenomic workflow for neoantigen prediction and selection. BMC Med Genomics. 2020;13:52. 10.1186/s12920-020-0683-4

48. Liu C, Zhang Y, Jian X, Tan X, Lu M, Ouyang J, et al. ProGeo-Neo v2.0: A One-Stop Software for Neoantigen Prediction and Filtering Based on the Proteogenomics Strategy. Genes (Basel). 2022;13:783. 10.3390/genes13050783

49. Wen B, Li K, Zhang Y, Zhang B. Cancer neoantigen prioritization through sensitive and reliable proteogenomics analysis. Nat Commun. 2020;11:1759. 10.1038/s41467-020-15456-w

50. Cox J, Mann M. MaxQuant enables high peptide identification rates, individualized p.p.b.-range mass accuracies and proteome-wide protein quantification. Nat Biotechnol. 2008;26:1367–72. 10.1038/nbt.1511

51. Huber F, Arnaud M, Stevenson BJ, Michaux J, Benedetti F, Thevenet J, et al. A comprehensive proteogenomic pipeline for neoantigen discovery to advance personalized cancer immunotherapy. Nat Biotechnol. 2024; 10.1038/s41587-024-02420-y

52. The Galaxy platform for accessible, reproducible and collaborative biomedical analyses: 2022 update. Nucleic Acids Res. 2022;50:W345–51. 10.1093/nar/gkac247

53. Galaxy Community. The Galaxy platform for accessible, reproducible, and collaborative data analyses: 2024 update. Nucleic Acids Res. 2024;52:W83–94. 10.1093/nar/gkae410

54. Galaxy Platform Directory: Servers, Clouds, and Deployable Resources - Galaxy Community Hub [Internet]. [cited 2025 Mar 24]. https://galaxyproject.org/use/. Accessed 24 Mar 2025

55. Mehta S, Bernt M, Chambers M, Fahrner M, Föll MC, Gruening B, et al. A Galaxy of informatics resources for MS-based proteomics. Expert Rev Proteomics. 2023;20:251–66. 10.1080/14789450.2023.2265062

56. Hiltemann S, Rasche H, Gladman S, Hotz H-R, Larivière D, Blankenberg D, et al. Galaxy Training: A powerful framework for teaching! PLoS Comput Biol. 2023;19:e1010752. 10.1371/journal.pcbi.1010752

57. Batut B, Hiltemann S, Bagnacani A, Baker D, Bhardwaj V, Blank C, et al. Community-Driven Data Analysis Training for Biology. Cell Syst. 2018;6:752–758.e1. 10.1016/j.cels.2018.05.012

58. Learning Pathway: Prediction of potential neoantigens [Internet]. [cited 2025 June 17]. https://training.galaxyproject.org/training-material/learning-pathways/neoantigen.html. Accessed 17 June 2025

59. Uhrig S, Ellermann J, Walther T, Burkhardt P, Fröhlich M, Hutter B, et al. Accurate and efficient detection of gene fusions from RNA sequencing data. Genome Res. 2021;31:448–60. 10.1101/gr.257246.119

60. Dobin A, Davis CA, Schlesinger F, Drenkow J, Zaleski C, Jha S, et al. STAR: ultrafast universal RNA-seq aligner. Bioinformatics. 2013;29:15–21. 10.1093/bioinformatics/bts635

61. Kim D, Paggi JM, Park C, Bennett C, Salzberg SL. Graph-based genome alignment and genotyping with HISAT2 and HISAT-genotype. Nat Biotechnol. 2019;37:907–15. 10.1038/s41587-019-0201-4

62. Wang X, Zhang B. customProDB: an R package to generate customized protein databases from RNA-Seq data for proteomics search. Bioinformatics. 2013;29:3235–7. 10.1093/bioinformatics/btt543

63. Pertea M, Pertea GM, Antonescu CM, Chang T-C, Mendell JT, Salzberg SL. StringTie enables improved reconstruction of a transcriptome from RNA-seq reads. Nat Biotechnol. 2015;33:290–5. 10.1038/nbt.3122

64. Pertea G, Pertea M. GFF Utilities: GffRead and GffCompare. F1000Res. 2020;9:ISCB Comm J-304. 10.12688/f1000research.23297.2

65. FragPipe [Internet]. FragPipe. [cited 2025 June 17]. https://fragpipe.nesvilab.org/. Accessed 17 June 2025

66. Yang KL, Yu F, Teo GC, Li K, Demichev V, Ralser M, et al. MSBooster: improving peptide identification rates using deep learning-based features. Nat Commun. 2023;14:4539. 10.1038/s41467-023-40129-9

67. Käll L, Canterbury JD, Weston J, Noble WS, MacCoss MJ. Semi-supervised learning for peptide identification from shotgun proteomics datasets. Nat Methods. 2007;4:923–5. 10.1038/nmeth1113

68. da Veiga Leprevost F, Haynes SE, Avtonomov DM, Chang H-Y, Shanmugam AK, Mellacheruvu D, et al. Philosopher: a versatile toolkit for shotgun proteomics data analysis. Nat Methods. 2020;17:869–70. 10.1038/s41592-020-0912-y

69. Johnson JE, Kumar P, Easterly C, Esler M, Mehta S, Eschenlauer AC, et al. Improve your Galaxy text life: The Query Tabular Tool. F1000Res. 2018;7:1604. 10.12688/f1000research.16450.2

70. Kong AT, Leprevost FV, Avtonomov DM, Mellacheruvu D, Nesvizhskii AI. MSFragger: ultrafast and comprehensive peptide identification in mass spectrometry-based proteomics. Nat Methods. 2017;14:513–20. 10.1038/nmeth.4256

71. Wen B, Zhang B. PepQuery2 democratizes public MS proteomics data for rapid peptide searching. Nat Commun. 2023;14:2213. 10.1038/s41467-023-37462-4

72. Nesvizhskii AI. Proteogenomics: concepts, applications and computational strategies. Nat Methods. 2014;11:1114–25. 10.1038/nmeth.3144

73. Altschul SF, Madden TL, Schäffer AA, Zhang J, Zhang Z, Miller W, et al. Gapped BLAST and PSI-BLAST: a new generation of protein database search programs. Nucleic Acids Res. 1997;25:3389–402. 10.1093/nar/25.17.3389

74. Kumar P, Johnson JE, McGowan T, Chambers MC, Heydarian M, Mehta S, et al. Discovering Novel Proteoforms Using Proteogenomic Workflows Within the Galaxy Bioinformatics Platform. Methods Mol Biol. 2025;2859:109–28. 10.1007/978-1-0716-4152-1_7

75. Robinson JT, Thorvaldsdóttir H, Winckler W, Guttman M, Lander ES, Getz G, et al. Integrative genomics viewer. Nat Biotechnol. 2011;29:24–6. 10.1038/nbt.1754

76. Raney BJ, Barber GP, Benet-Pagès A, Casper J, Clawson H, Cline MS, et al. The UCSC Genome Browser database: 2024 update. Nucleic Acids Res. 2024;52:D1082–8. 10.1093/nar/gkad987

77. Carithers LJ, Moore HM. The Genotype-Tissue Expression (GTEx) Project. Biopreserv Biobank. 2015;13:307–8. 10.1089/bio.2015.29031.hmm

78. Krysiak K, Danos AM, Kiwala S, McMichael JF, Coffman AC, Barnell EK, et al. A community approach to the cancer-variant-interpretation bottleneck. Nat Cancer. 2022;3:522–5. 10.1038/s43018-022-00379-w

79. Tomczak K, Czerwińska P, Wiznerowicz M. The Cancer Genome Atlas (TCGA): an immeasurable source of knowledge. Contemp Oncol (Pozn). 2015;19:A68–77. 10.5114/wo.2014.47136

80. Desiere F, Deutsch EW, King NL, Nesvizhskii AI, Mallick P, Eng J, et al. The PeptideAtlas project. Nucleic Acids Res. 2006;34:D655–658. 10.1093/nar/gkj040

81. Lu M, Xu L, Jian X, Tan X, Zhao J, Liu Z, et al. dbPepNeo2.0: A Database for Human Tumor Neoantigen Peptides From Mass Spectrometry and TCR Recognition. Front Immunol. 2022;13:855976. 10.3389/fimmu.2022.855976

82. Shao Y, Ge S, Dong R, Ji W, Qin C, Wen P. NeoTImmuML: a machine learning-based prediction model for human tumor neoantigen immunogenicity. Front Immunol. 2025;16:1681396. 10.3389/fimmu.2025.1681396

83. Szolek A, Schubert B, Mohr C, Sturm M, Feldhahn M, Kohlbacher O. OptiType: precision HLA typing from next-generation sequencing data. Bioinformatics. 2014;30:3310–6. 10.1093/bioinformatics/btu548

84. Boegel S, Bukur T, Castle JC, Sahin U. In Silico Typing of Classical and Non-classical HLA Alleles from Standard RNA-Seq Reads. Methods Mol Biol. 2018;1802:177–91. 10.1007/978-1-4939-8546-3_12

85. Jurtz V, Paul S, Andreatta M, Marcatili P, Peters B, Nielsen M. NetMHCpan-4.0: Improved Peptide-MHC Class I Interaction Predictions Integrating Eluted Ligand and Peptide Binding Affinity Data. J Immunol. 2017;199:3360–8. 10.4049/jimmunol.1700893

86. Pagliuca S, Gurnari C, Rubio MT, Visconte V, Lenz TL. Individual HLA heterogeneity and its implications for cellular immune evasion in cancer and beyond. Front Immunol. 2022;13:944872. 10.3389/fimmu.2022.944872

87. pVACbind — pVACtools 6.0.3 documentation [Internet]. [cited 2025 Dec 2]. https://pvactools.readthedocs.io/en/latest/pvacbind.html. Accessed 2 Dec 2025

88. Jagtap PD, Johnson JE, Onsongo G, Sadler FW, Murray K, Wang Y, et al. Flexible and accessible workflows for improved proteogenomic analysis using the Galaxy framework. J Proteome Res. 2014;13:5898–908. 10.1021/pr500812t

89. Pandey K, Mifsud NA, Lim Kam Sian TCC, Ayala R, Ternette N, Ramarathinam SH, et al. In-depth mining of the immunopeptidome of an acute myeloid leukemia cell line using complementary ligand enrichment and data acquisition strategies. Mol Immunol. 2020;123:7–17. 10.1016/j.molimm.2020.04.008

90. Chong C, Müller M, Pak H, Harnett D, Huber F, Grun D, et al. Integrated proteogenomic deep sequencing and analytics accurately identify non-canonical peptides in tumor immunopeptidomes. Nat Commun. 2020;11:1293. 10.1038/s41467-020-14968-9

91. Trujillo JA, Croft NP, Dudek NL, Channappanavar R, Theodossis A, Webb AI, et al. The cellular redox environment alters antigen presentation. J Biol Chem. 2014;289:27979–91. 10.1074/jbc.M114.573402

92. Chen Z, He X. Application of third-generation sequencing in cancer research. Med Rev (2021). 2021;1:150–71. 10.1515/mr-2021-0013

93. Blankenberg D, Von Kuster G, Coraor N, Ananda G, Lazarus R, Mangan M, et al. Galaxy: a web-based genome analysis tool for experimentalists. Curr Protoc Mol Biol. 2010;Chapter 19:Unit 19.10.1-21. 10.1002/0471142727.mb1910s89

94. Xia H, Hoang MH, Schmidt E, Kiwala S, McMichael J, Skidmore ZL, et al. pVACview: an interactive visualization tool for efficient neoantigen prioritization and selection. Genome Med. 2024;16:132. 10.1186/s13073-024-01384-7

95. Pyke RM, Mellacheruvu D, Dea S, Abbott C, Zhang SV, Phillips NA, et al. Precision Neoantigen Discovery Using Large-Scale Immunopeptidomes and Composite Modeling of MHC Peptide Presentation. Mol Cell Proteomics. 2023;22:100506. 10.1016/j.mcpro.2023.100506

96. Xu L, Yang Q, Dong W, Li X, Wang K, Dong S, et al. Meta learning for mutant HLA class I epitope immunogenicity prediction to accelerate cancer clinical immunotherapy. Brief Bioinform. 2024;26:bbae625. 10.1093/bib/bbae625

97. Lippolis JD, White FM, Marto JA, Luckey CJ, Bullock TNJ, Shabanowitz J, et al. Analysis of MHC class II antigen processing by quantitation of peptides that constitute nested sets. J Immunol. 2002;169:5089–97. 10.4049/jimmunol.169.9.5089

98. diaPASEF Analysis for HLA-I Peptides Enables Quantification of Common Cancer Neoantigens - PubMed [Internet]. [cited 2025 June 17]. https://pubmed.ncbi.nlm.nih.gov/40044040/. Accessed 17 June 2025

99. Sensitive Immunopeptidomics by Leveraging Available Large-Scale Multi-HLA Spectral Libraries, Data-Independent Acquisition, and MS/MS Prediction - PubMed [Internet]. [cited 2025 June 17]. https://pubmed.ncbi.nlm.nih.gov/33845167/. Accessed 17 June 2025

100. Rajczewski AT, Blakeley-Ruiz JA, Meyer A, Vintila S, McIlvin MR, Van Den Bossche T, et al. Data-Independent Acquisition Mass Spectrometry as a Tool for Metaproteomics: Interlaboratory Comparison Using a Model Microbiome. Proteomics. 2025;25:e202400187. 10.1002/pmic.202400187

101. Li K, Teo GC, Yang KL, Yu F, Nesvizhskii AI. diaTracer enables spectrum-centric analysis of diaPASEF proteomics data. Nat Commun. 2025;16:95. 10.1038/s41467-024-55448-8

102. Yu F, Teo GC, Kong AT, Fröhlich K, Li GX, Demichev V, et al. Analysis of DIA proteomics data using MSFragger-DIA and FragPipe computational platform. Nat Commun. 2023;14:4154. 10.1038/s41467-023-39869-5

103. Karunratanakul K, Tang H-Y, Speicher DW, Chuangsuwanich E, Sriswasdi S. Uncovering Thousands of New Peptides with Sequence-Mask-Search Hybrid De Novo Peptide Sequencing Framework. Mol Cell Proteomics. 2019;18:2478–91. 10.1074/mcp.TIR119.001656

104. Galaxy [Internet]. [cited 2025 June 17]. https://usegalaxy.eu/. Accessed 17 June 2025

105. Diop D, Chauvin C, Jean-Jean O. Aminoglycosides and other factors promoting stop codon readthrough in human cells. C R Biol. 2007;330:71–9. 10.1016/j.crvi.2006.09.001

106. Lim Kam Sian TCC, Goncalves G, Steele JR, Shamekhi T, Bramberger L, Jin D, et al. SAPrIm, a semi-automated protocol for mid-throughput immunopeptidomics. Front Immunol. 2023;14:1107576. 10.3389/fimmu.2023.1107576

107. Pandey K, Ramarathinam SH, Purcell AW. Isolation of HLA Bound Peptides by Immunoaffinity Capture and Identification by Mass Spectrometry. Curr Protoc. 2021;1:e92. 10.1002/cpz1.92

108. freebayes/freebayes: Bayesian haplotype-based genetic polymorphism discovery and genotyping. [Internet]. [cited 2025 June 17]. https://github.com/freebayes/freebayes. Accessed 17 June 2025

109. Bowler-Barnett EH, Fan J, Luo J, Magrane M, Martin MJ, Orchard S, et al. UniProt and Mass Spectrometry-Based Proteomics-A 2-Way Working Relationship. Mol Cell Proteomics. 2023;22:100591. 10.1016/j.mcpro.2023.100591

110. Adusumilli R, Mallick P. Data Conversion with ProteoWizard msConvert. Methods Mol Biol. 2017;1550:339–68. 10.1007/978-1-4939-6747-6_23

111. Galaxy | Tool Shed [Internet]. [cited 2025 Mar 24]. https://toolshed.g2.bx.psu.edu/. Accessed 24 Mar 2025

112. FragPipe/LICENSE at develop · Nesvilab/FragPipe [Internet]. GitHub. [cited 2025 Mar 24]. https://github.com/Nesvilab/FragPipe/blob/develop/LICENSE. Accessed 24 Mar 2025

113. galaxy/LICENSE.txt at dev · galaxyproject/galaxy [Internet]. GitHub. [cited 2025 Mar 24]. https://github.com/galaxyproject/galaxy/blob/dev/LICENSE.txt. Accessed 24 Mar 2025

