## Supplemental Figures and Table for "A modular, immunopeptidogenomic (iPepGen) analysis pipeline for discovery, verification, and prioritization of cancer peptide neoantigen candidates"

**Supplemental Figure 1:** Venn Diagram comparing peptide identifications and overlap between FragPipe (red) and PEAKS (green). Database search settings were identical (no enzyme specificity, methionine oxidation as variable modification) using the same raw immunopeptidomics MS/MS file and customized protein sequence database of 2.9M+ sequences from MPNST cells as described in the main text, with both reporting results at 1% peptide FDR.

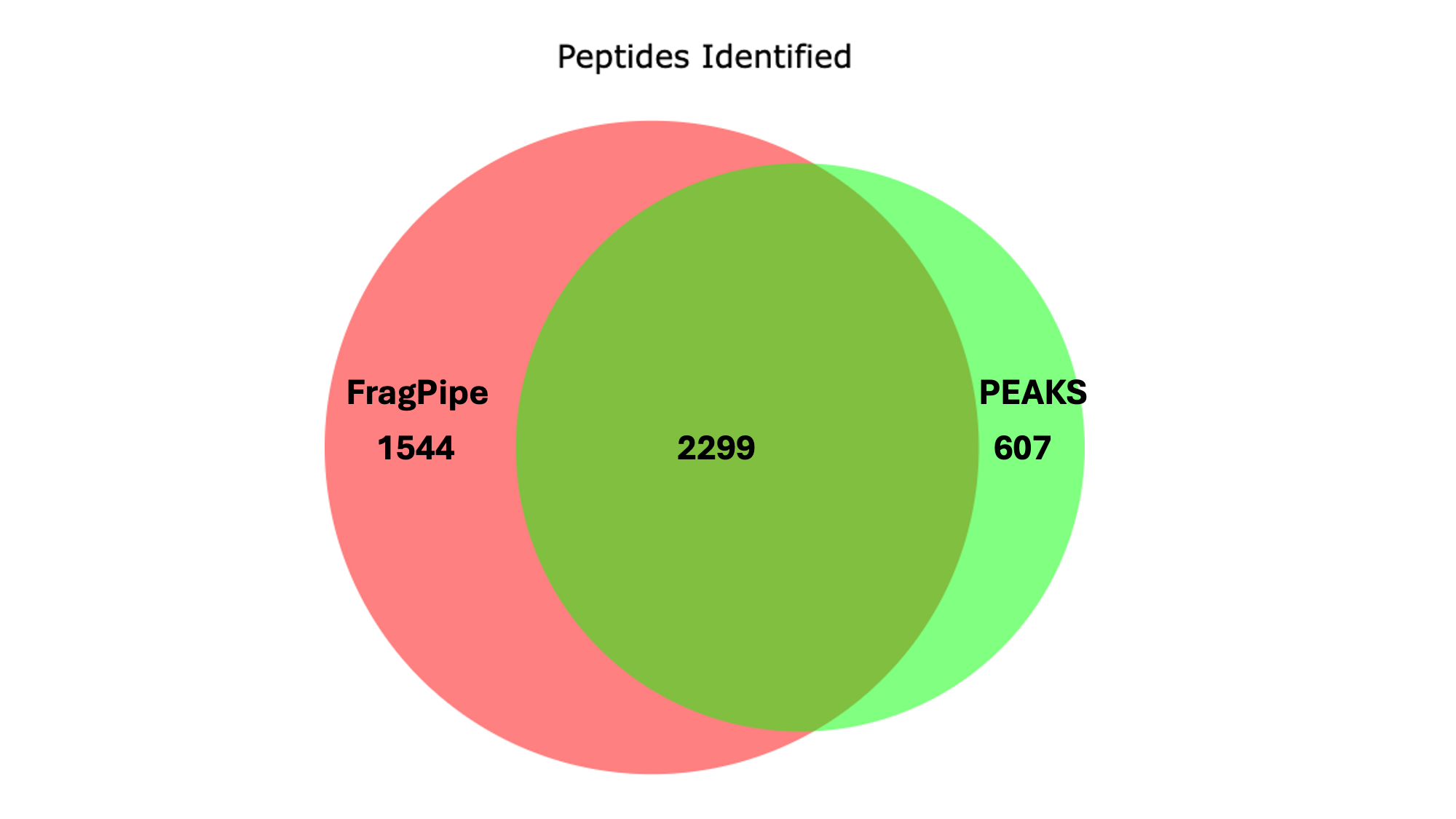

**Supplemental Figure 2:** Comparison of peptide identifications with and without MSBooster. Bar plots show the mean number of peptides identified per replicate across two peptide categories: **All Peptides** and **Non-Reference Peptides**. Results are displayed for runs processed **with MSBooster** (red) and **without MSBooster** (teal). Three technical replicate raw MS/MS immunopeptidomics files were analyzed from the published study by the Purcell group (Curr Protoc. 2021;1:e92). Error bars represent standard deviations across replicates. A broken y-axis is used to display the large difference in scale between total peptide identifications and the smaller counts for non-reference peptides

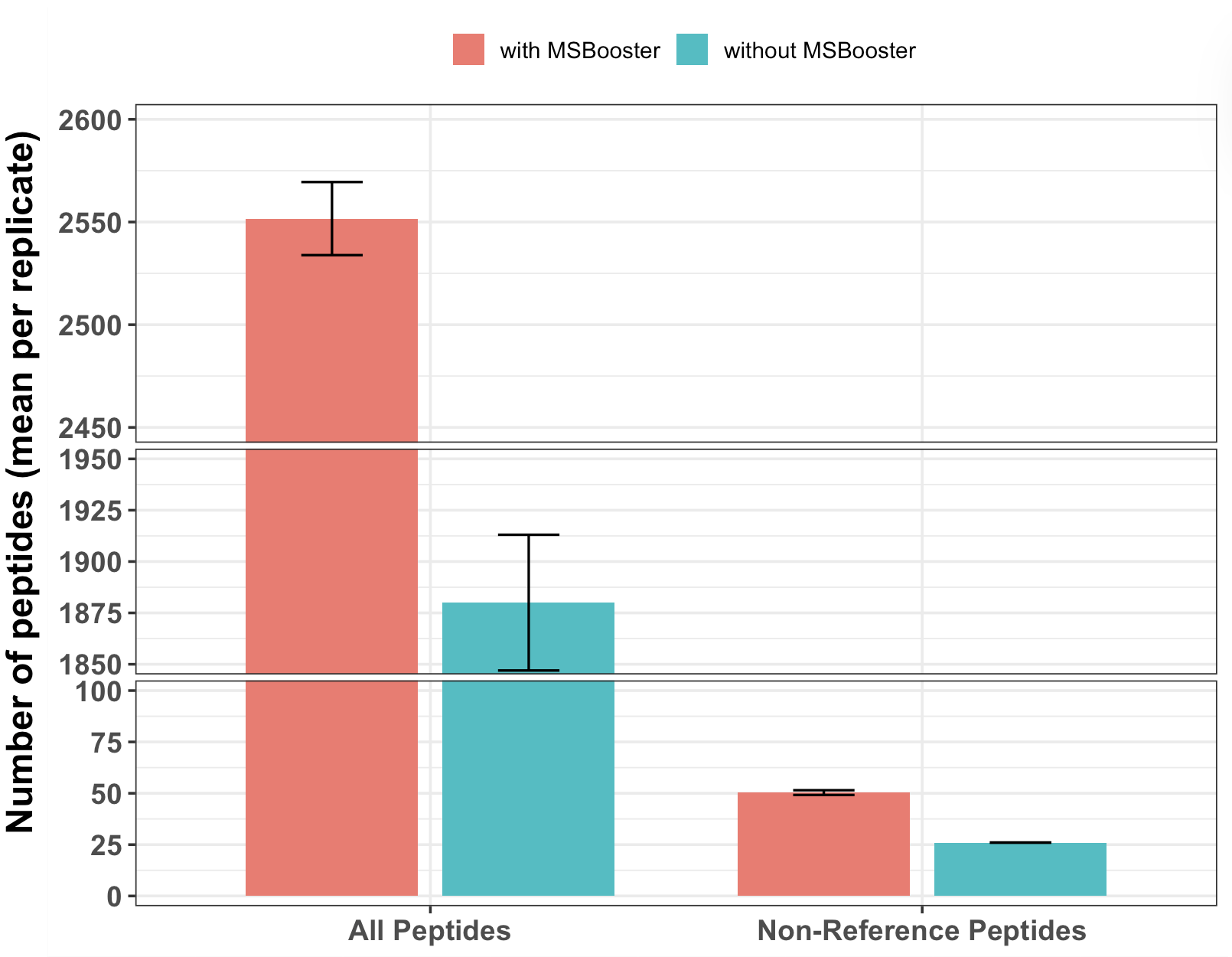

**Supplemental Figure 3:** Funnel diagram illustrating the multi-step peptide filtering workflow used to identify high-confidence novel peptides from MS/MS data. Starting with approximately 2.9 million predicted sequences and one RAW MS/MS dataset, FragPipe initially detected 3,843 peptides. Of these, 123 were classified as non-reference peptides (not present in the canonical proteome). PepQuery validation further reduced the set to 28 high-confidence peptide-spectrum matches. Final BLAST-P filtering removed peptides with homology to known proteins, yielding 19 novel peptide candidates that passed all validation criteria.

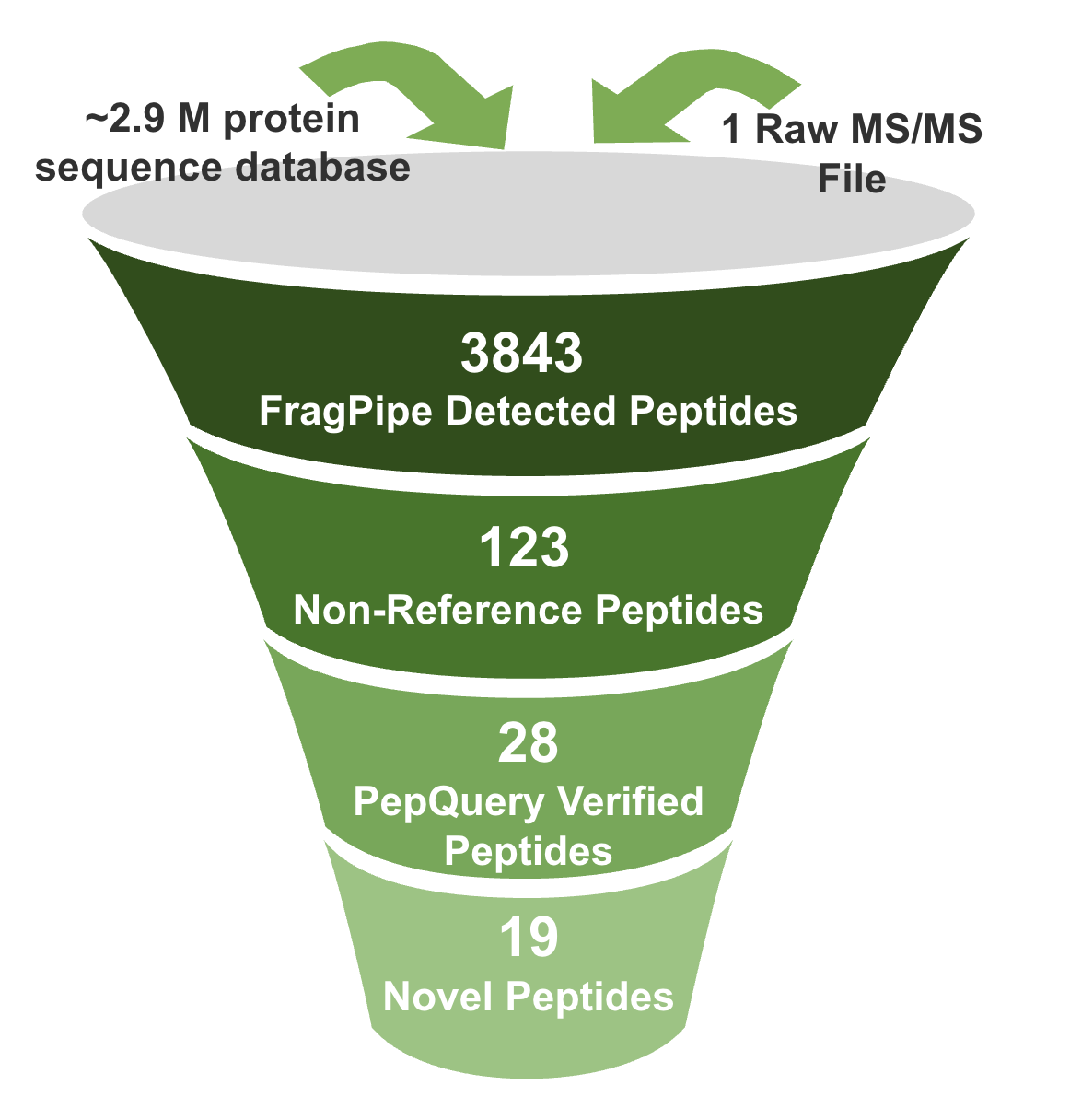

**Supplemental Figure 4:** Visualization of a neoantigen candidate in the UCSC browser. The PepPointer tool in Module 4 provides a URL that can be inputted directly to the UCSC browser. Among many other functionalities, the browser allows for visualization of transcripts encoding the peptide across cancer tissues (from the TCGA resource) and expression in normal tissues from the GTEx database. The example shown is for the neoantigen peptide RLMGEGTSSL, identified in the demonstration data from the STS-26T MPNST cells, which contained amino acids encoded by a portion of 5’-UTR sequence for the gene ARHGAP11B. Mousing over the bar chart images in each track provides information on transcript expression levels in specific tissues.

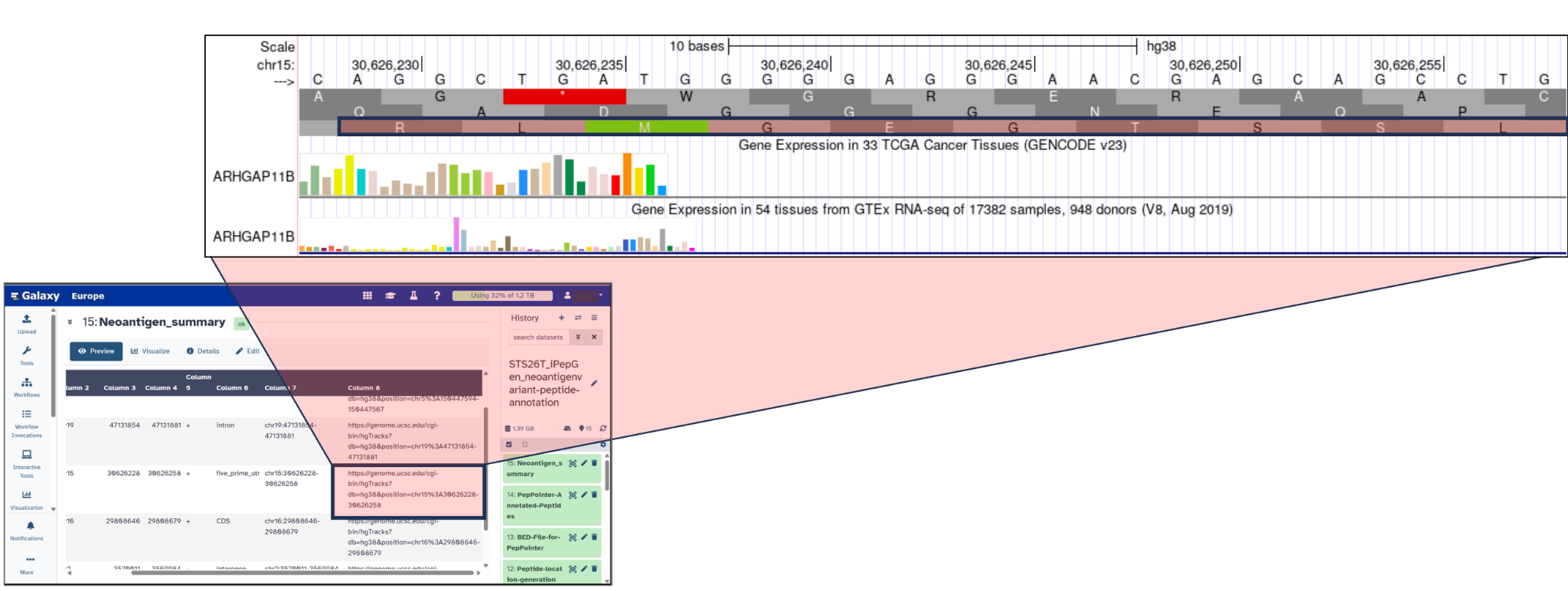

**Supplemental Figure 5. Manual characterization of the genomic mapping of a complex transcript coding an identified neoantigen candidate** **(A)** PepPointer analysis annotates the peptide (sequence HSEVQTLKY, **red highlight**) as being located within an intronic region. **(B)** Genomic alignment indicates that the peptide is absent from the predicted genomic location. **(C)** BLAST analysis confirms the peptide’s alignment to a known protein, supporting its chromosomal location, sequence alignment with an amino acid substitution in the SKA3 protein. **(D)** A re-analysis using IGV with a three-frame translation (**red highlight**) demonstrates that eight of the nine amino acids composing the peptide are coded from a transcript mapping to the SKA3 gene structure; however the transcript also contains additional novel sequence mapping to an intron and adding amino acid, making this a novel non-reference neoantigen candidate coded by a combination of exon and intron sequences. Such cases are novel and not easily annotated by the PepPointer tool and require manual exploration.

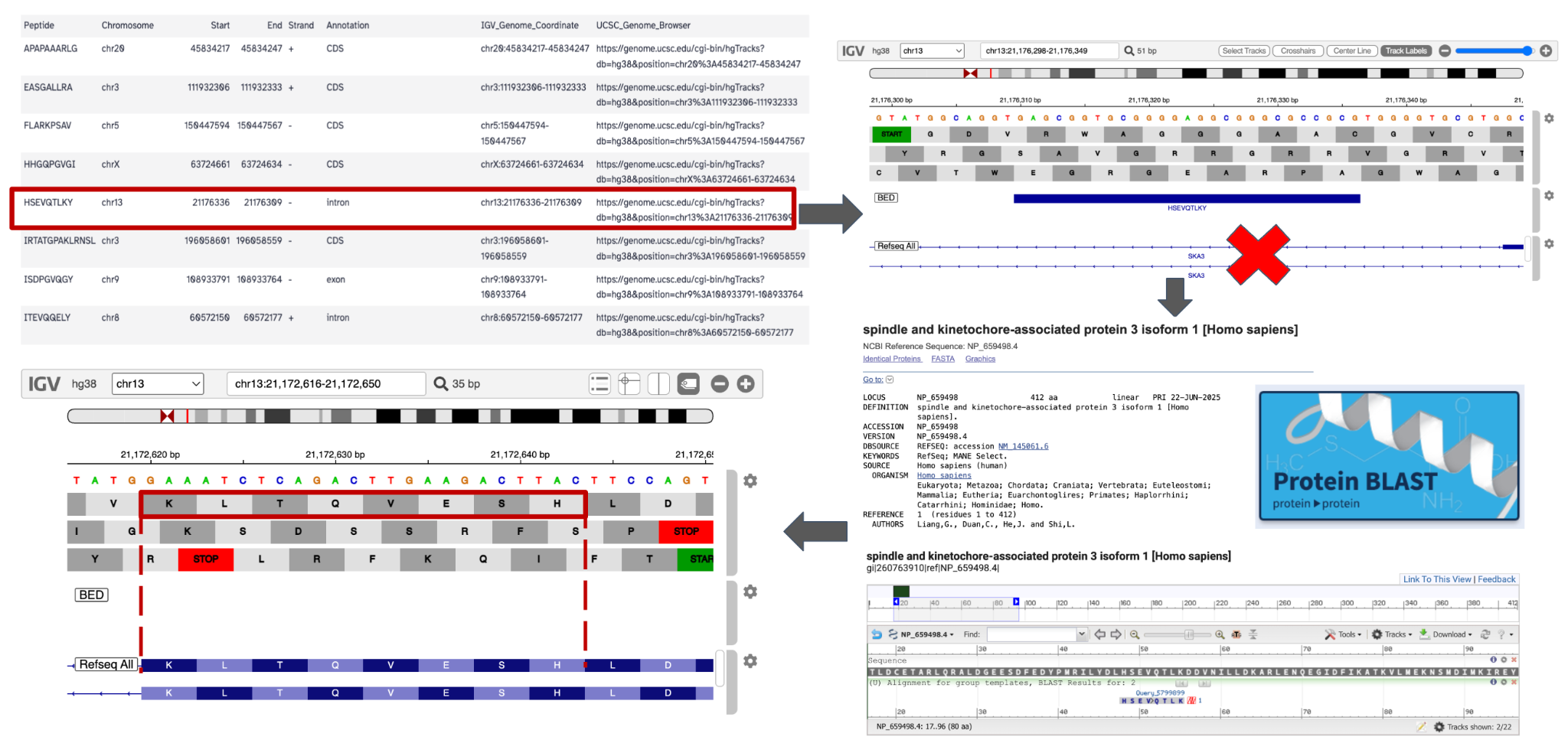

**Supplemental Figure 6A. Preparing FASTA peptide list input for pVACbind**

Search for and select “Filter FASTA” from the tool panel on the left side of the Galaxy GUI. In the Filter FASTA tool select these settings:

For the **FASTA sequences**, select “IEDB PEPTIDES FASTA” item from the iPepGen analysis history

For **Criteria for filtering on the headers** select “Regular expression on the headers” from the dropdown menu

In the input field **Regular expression pattern the header should match** enter the following expression: **^>.***

In the **Criteria for filtering on the sequences** select “No filtering” from the dropdown menu

Toggle **Remove duplicate sequences** to “Yes”

Click on “**Run tool**”

A new history item containing the filtered FASTA file with redundant entries removed will be created.

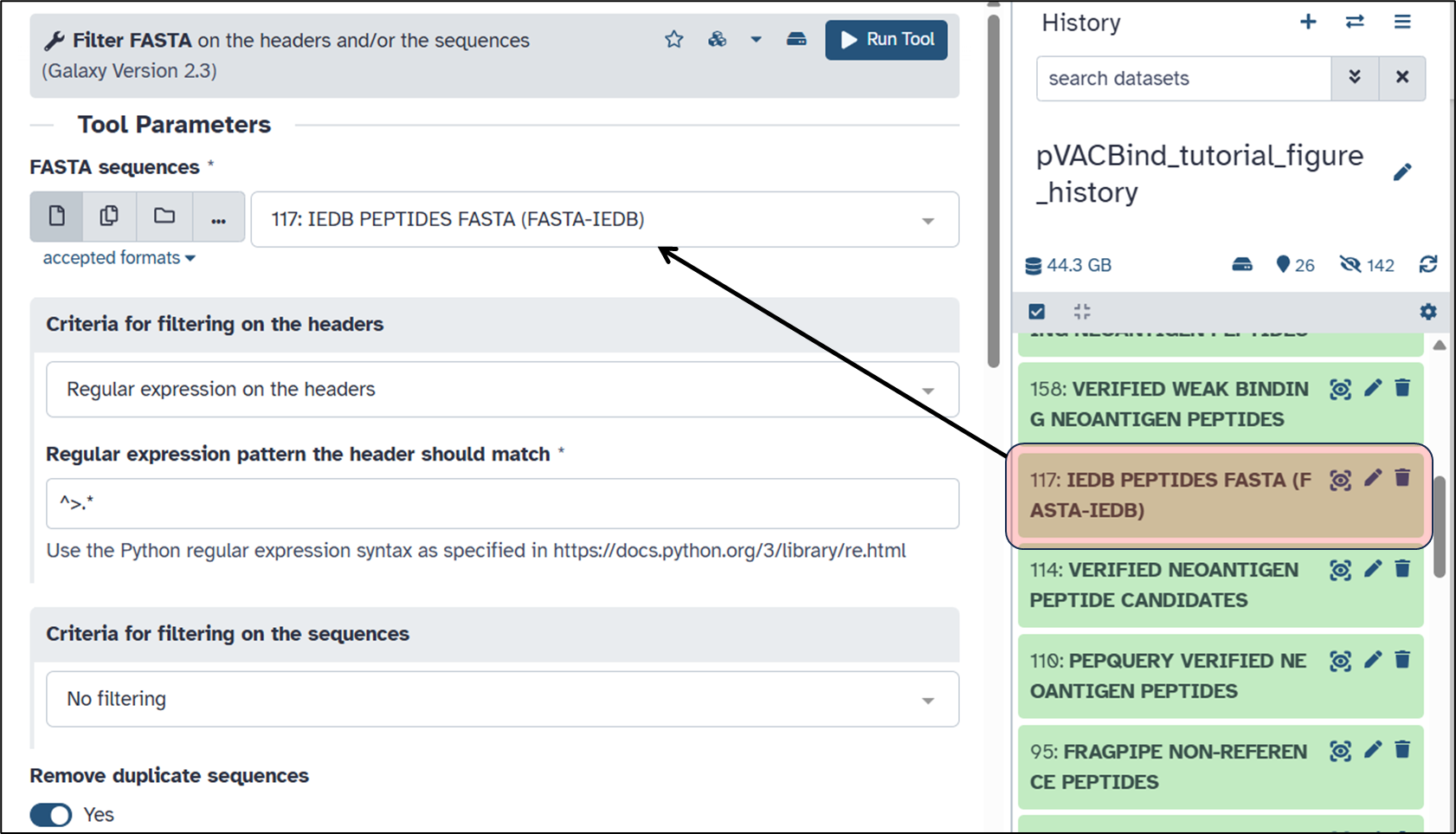

**Supplemental Figure 6b. Running pVACbind.**

Search for and select “pVACbind” from the tool panel on the left side of the Galaxy GUI. In the pVACbind tool settings select:

For **Alleles** select “From history” in the dropdown menu

For **Alleles File** select “COMBINED OPTITYPE AND SEQ2HLA HLA GENOTYPES” from the iPepGen history (the search bar in the history column can be used to find this)

For **Input File** select the newly created, non-redundant filtered peptide FASTA file in the active history from step 1 above

For **Sample Name**, input any text identifier desired for the current analysis

For **Algorithm**, select any combination of algorithms from the dropdown menu; for the example shown here algorithms for MHC class I analysis were selected

For **Class I Epitope Length** and **Class II Epitope Length** select desired length of peptides to consider for analysis, based on the length range specified for Module 3 verification; at least one peptide length is needed in the both the class I and class II fields, even if only conducting a single class analysis

For all other parameters leave as “default” settings as a starting point (see pVACbind documentation for changing other tool settings at https://pvactools.readthedocs.io/en/latest/pvacbind.html)

Click on **“Run tool”**

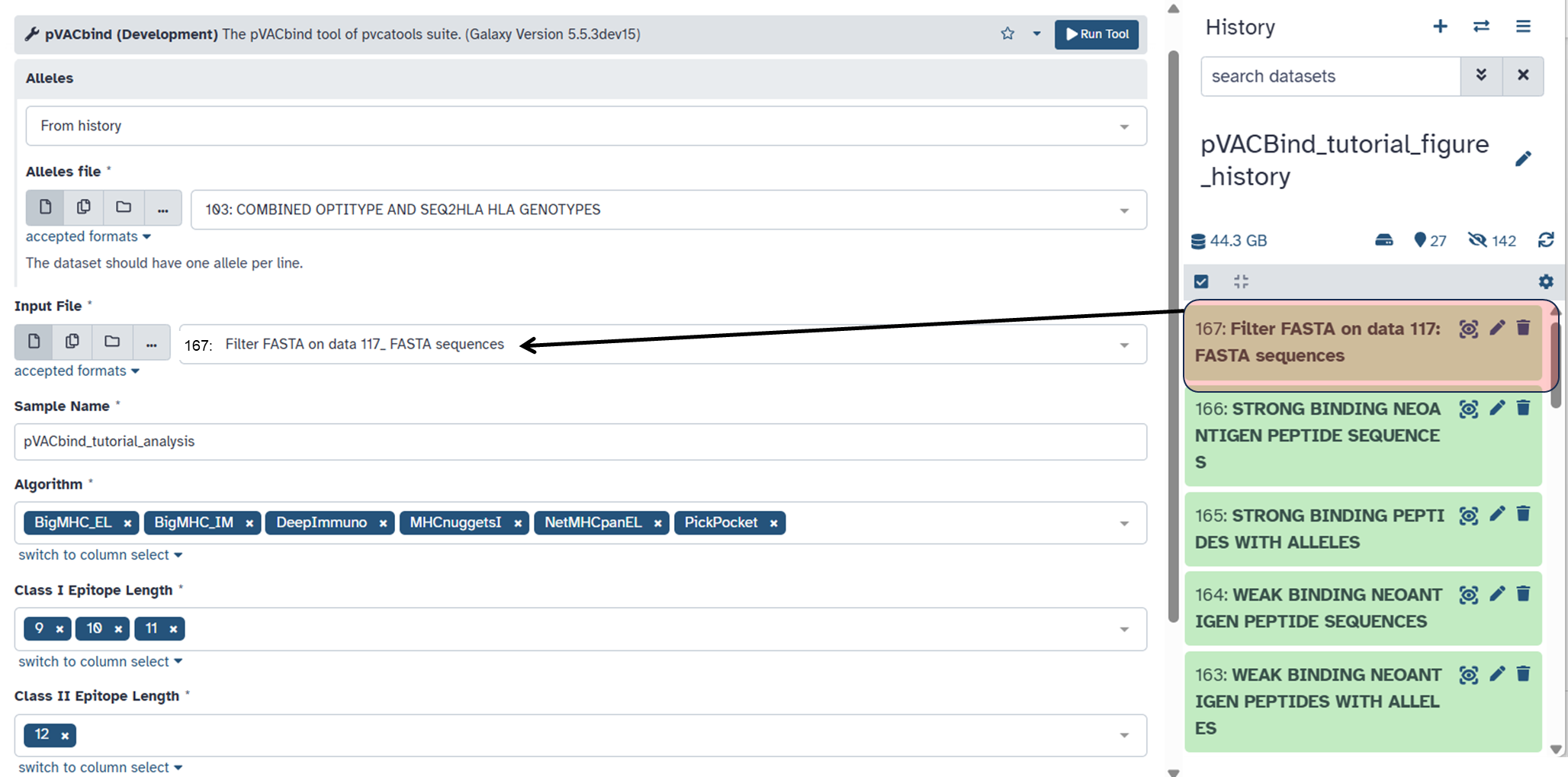

**Supplemental Figure 6C. Working with outputs from pVACbind.**

A collection of MHC Class I outputs and/or MHC Class II will be output; if only tools for analysis of one class were selected, one of these outputs will be blank. In the example below, only MHC Class I tools were used, so the Class II Output collection contains no datasets.

The collection of outputs contains several tabular files, including:

“pVACbind_tutorial_analysis.all_epitopes.tsv”: A tabular file with results for all of the allele peptide pairs analyzed, complete with scoring from selected algorithms

“pVACbind_tutorial_analysis.filtered.tsv”: This tabular file shows the best binding and/or immunogenicity scores for each peptide evaluated, along with the HLA allele analyzed to generate these scores. See <https://pvactools.readthedocs.io/en/latest/pvacbind.html> for more information on these outputs and filtering methods.

The center panel in the figure below shows an example of the filtered tabular output from pVACbind.

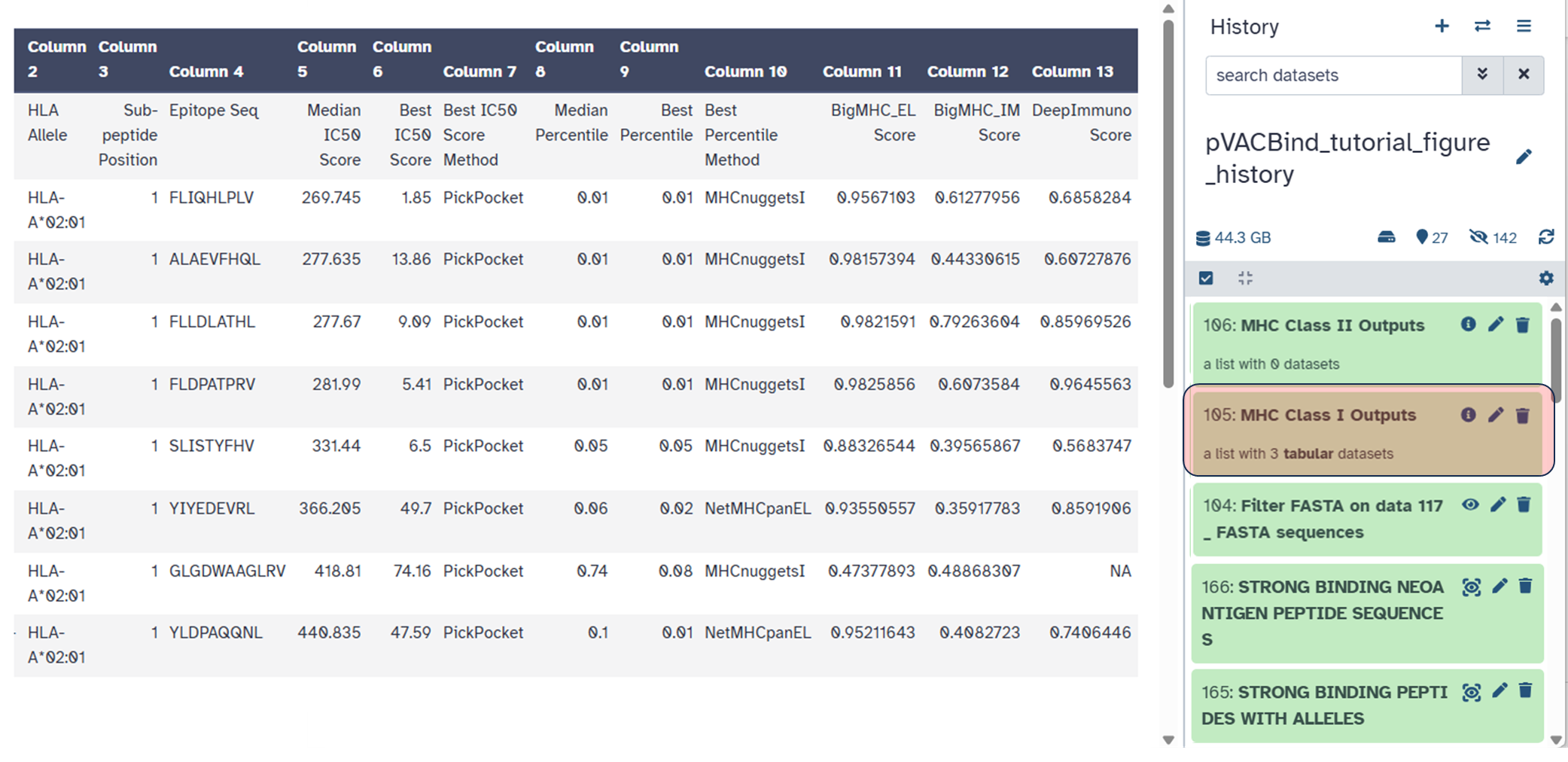

**Supplemental Figure 7:** Example peptides identified and quantified by LFQ via the customized workflow integrating modules 2 and 3 of the iPepGen pipeline. Eight separate immunopeptidomics raw files were analyzed from a previously published study (*Nat Commun.* 2020 11:1293), with four of these being data from control melanoma cells and four from melanoma cells treated with 5-aza-2′-deoxycytidine (DAC), which promotes antigen peptide antigen presentation on the HLA complex. **A)** A neoantigen peptide candidate that was discovered and verified by the workflow matching a non-reference sequence and showing a significant increase abundance in the DAC-treated samples as measured by LFQ intensities.  **B)** A TAA peptide belongs to the melanoma-associated antigen 8/9 protein (MAGE8/9), which is a known antigen in melanoma cells and shows an expected increase in abundance with DAC treatment.

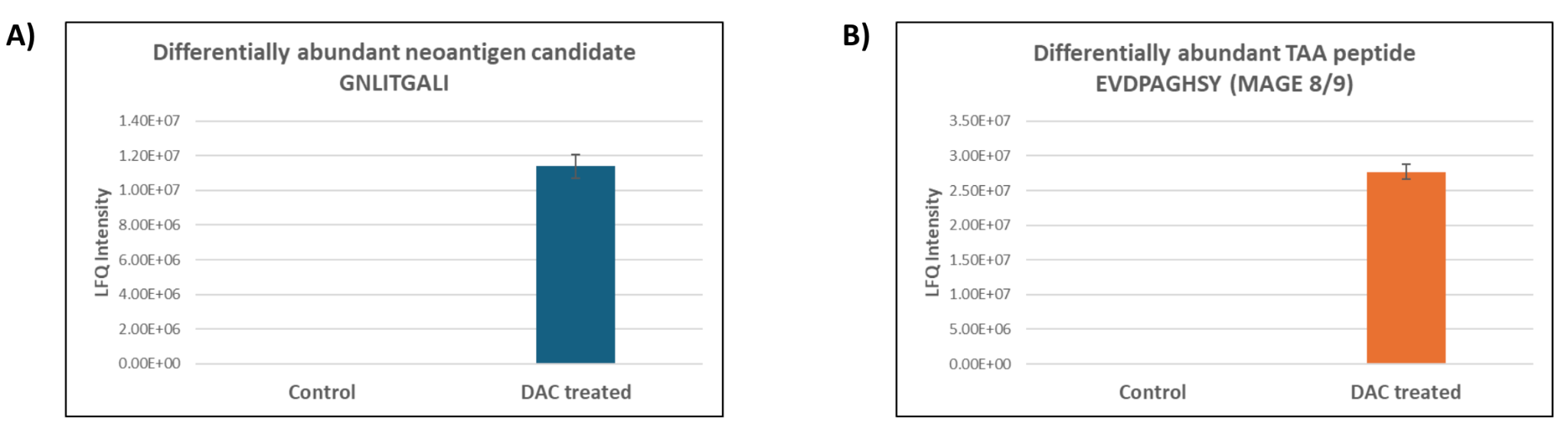

**Supplemental Table 1.** Runtime and Memory Benchmarking of FragPipe (Version 23.0) Across Databases of Different Sizes

| **Database Size** | **Number of Candidate Sequences** | **Average Runtime** | **Average Memory Usage (GB)** | **Processor Cores Used** |
| --- | --- | --- | --- | --- |
| Small | 100,000 | 29 minutes | 29.0 | 12 |
| Medium | 1,000,000 | 43 minutes | 127.3 | 12 |
| Original | 2,907,272 | 1 hour 36 minutes | 256.0 | 12 |
